# CRYO-EM STRUCTURE OF THE DELTA-RETROVIRAL INTASOME IN COMPLEX WITH THE PP2A REGULATORY SUBUNIT B56γ

**DOI:** 10.1101/2020.06.19.161513

**Authors:** Michał S. Barski, Jordan J. Minnell, Zuzana Hodakova, Valerie E. Pye, Andrea Nans, Peter Cherepanov, Goedele N. Maertens

## Abstract

The *Retroviridae* delta-retrovirus genus includes the most oncogenic pathogen– human T-cell lymphotropic virus type 1 (HTLV-1)(1). Many of the ~20 million people infected with HTLV-1 will develop severe leukaemia (2) or an ALS-like motor disease (3) unless a therapy becomes available. A key step in the establishment of infection is the integration of viral genetic material into the host genome, catalysed by the viral integrase (IN) enzyme. Here we used X-ray crystallography and single-particle cryo-electron microscopy to determine the structure of functional delta-retroviral IN assembled on viral DNA ends and bound the B56γ subunit of its human host factor, the protein phosphatase 2A (4). The structure reveals a tetrameric IN assembly bound to the phosphatase via a conserved short linear motif found within the extended linker connecting the catalytic core (CCD) and C-terminal (CTD) IN domains. Unexpectedly, all four IN subunits are involved in B56γ binding, taking advantage of the flexibility of the CCD-CTD linkers. Our results fill the current gap in the structural understanding of the delta-retroviral integration machinery. Insight into the interactions between the delta-retroviral intasome and the host will be crucial for understanding the pattern of integration events in infected individuals and therefore bears important clinical implications.

---

Despite the severe pathology caused by HTLV-1, including adult T-cell leukaemia/lymphoma (ATLL) (2), myelopathy (HAM/TSP) (3) and uveitis (5), most aspects of delta-retrovirus biochemistry remain *terra incognita*. Elucidation of mechanisms behind delta-retroviral integration is particularly important in order to address issues of much needed pharmacological intervention, and to understand how integration site targeting affects clonal expansion of malignant T-cells leading to ATLL.

Reverse transcription of retroviral RNA yields linear double stranded viral DNA (vDNA) with a copy of long terminal repeat (LTR) at each end. Integrase (IN) binds and brings together the vDNA ends in the intasome nucleoprotein complex to insert them into host chromosomal DNA (6). IN first catalyses 3'-processing reaction of the vDNA ends, exposing reactive 3' OH nucleophile groups that it uses to attack a pair of phosphodiester bonds within target DNA (tDNA), resulting in strand transfer. After host-mediated DNA repair, the provirus becomes permanently integrated into the host chromatin. Although the structural mechanics of this reaction was revealed almost a decade ago with the example of a spumaviral intasome (7–9), the architectures of the intasome nucleoprotein complexes exhibit remarkable variety among the retroviral genera: ranging from tetrameric (spumaviruses) (7), through octameric (betaretrovirus (10) and alpharetrovirus (11)) to dodeca/hexadecameric (lentiviruses)(12,13) assemblies. The functional organization of IN in the delta-retroviral intasome is unknown. An added layer of complexity is the recruitment of IN-binding host proteins, which in many cases help to guide integration to specific genomic loci {Hare, 2009 #372;Maertens, 2016 #46;Sharma, 2013 #117;De Rijck, 2013 #358;Gupta, 2013 #370}. The structural characterisation of such virus-host complexes is undetermined.

We found that IN from simian T-lymphotropic virus type 1 (STLV-1), which shares 83% amino acid sequence identity with its counterpart from HTLV-1 (Figure S1), is competent for concerted strand transfer activity *in vitro*. Akin to HTLV-1 IN, the enzyme readily utilizes short, double-stranded oligonucleotide mimics of vDNA ends (Figure S2A-B) (4,18) for integration. Formation of stable intasomes *in vitro* can be technically challenging, and often requires the presence of host factors and/or hyperactivating mutations (12,13,19). In one approach, the positively charged N-terminal region of LEDGF was fused with HIV-1 IN, in order to promote formation of intasomes *in vitro* (20). The delta-retroviral IN host cofactor B56γ (4), as part of the heterotrimeric protein phosphatase 2A (PP2A) holo-enzyme, interacts with a number of chromatin-associated proteins (21–24) and potently stimulates concerted integration activity of delta-retroviral INs (4). To aid stable intasome formation without altering IN, we constructed a LEDGF/ΔIBD-B56γ chimera containing the DNA-binding region of LEDGF (residues 1-324) and B56γ (residues 11-380) (Figure S2C). Electrophoretic mobility shift assays (EMSAs) showed that STLV-1 IN forms a stable nucleoprotein complex with vDNA in the presence of LEDGF/ΔIBD-B56γ (Figure S2D). Size-exclusion chromatography of the assembly reactions revealed a high-molecular weight species that was competent for strand transfer activity (Figure S3A and B). Negative-stain electron microscopy (EM) analysis of peak fractions identified distinct particles measuring ~15 nm in the longest dimension, with prominent two-fold symmetry (Figure S3C-E).

To characterize the delta-retroviral intasome at near-atomic resolution, we imaged the particles by cryogenic electron microscopy (cryoEM). The nucleoprotein samples were vitrified on open hole grids as well as absorbed onto graphene oxide film, resulting in two anisotropically sampled yet complementary datasets. Merging the data allowed us to refine an isotropic 3D reconstruction to an overall resolution of 3.37 Å, and 2.9 Å throughout the conserved intasome core (CIC) region (Figures S4 – S6, and S7A). Due to the lack of structural information on delta-retroviral INs, we determined a number of crystal structures spanning the catalytic core domain (CCD) (Figures S8 and Table S3) and the C-terminal domain (CTD) of HTLV-2 and −1 IN, respectively, to aid the interpretation of the cryoEM map. The IN/CTD structure was determined in isolation as well as in complex with B56γ (Figure S10 - S11 and Table S4). The apo CTD crystal structure shows a canonical, small β-barrel SH3-like fold, with side-to-side orientation similar to that previously seen in a HIV-1 IN/CTD crystal structure (PDB ID 5TC2) (Figure S10C and D). The short linear motif (SLiM) harboured by IN within the CCD-CTD linker is clearly resolved, bound to a groove in the centre of B56γ; the previously characterised binding site for endogenous substrates of PP2A (Figure 2 and Figure S11)(25,26).

**Figure 1.**
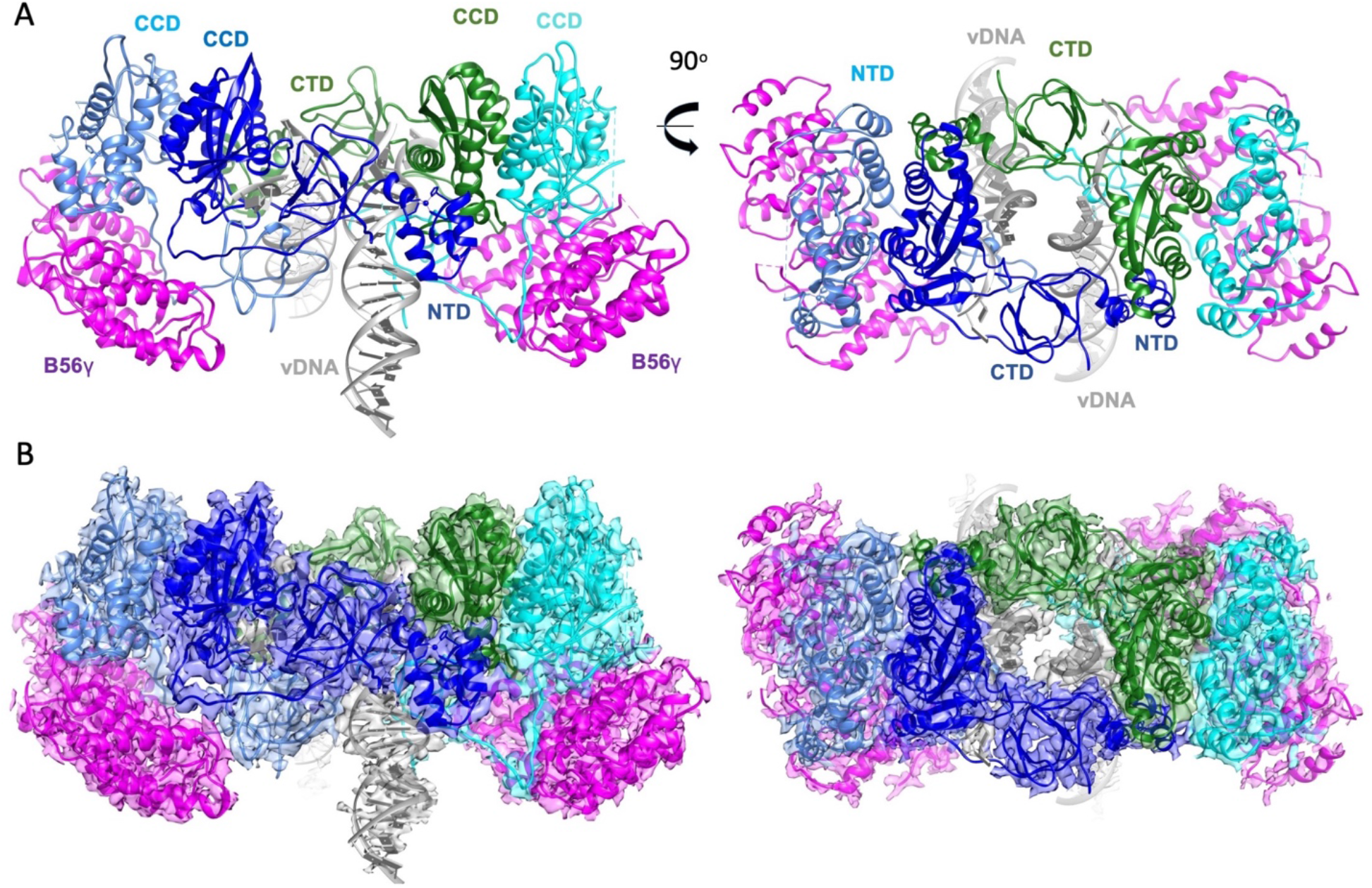
Structure of the delta-retroviral intasome in complex with the B56γ regulatory subunit of PP2A. **A**, The intasome-B56γ structure. Delta-retroviral intasome attains a tetrameric arrangement - IN chains are coloured in dark and light blue, their symmetrical counterparts in cyan and green. Viral DNA (vDNA) in grey is seen in the centre, complexed by extensive interactions with all IN domains. B56γ (pink) flanks either side of the intasome, targeting all four available IN CCD-CTD linkers. Both side view (left) as well as top-down view (right) are shown. **B**, 3.37 Å cryoEM map reconstruction is overlaid on top of the model in orientations analogous to panel A.

**Figure 2.**
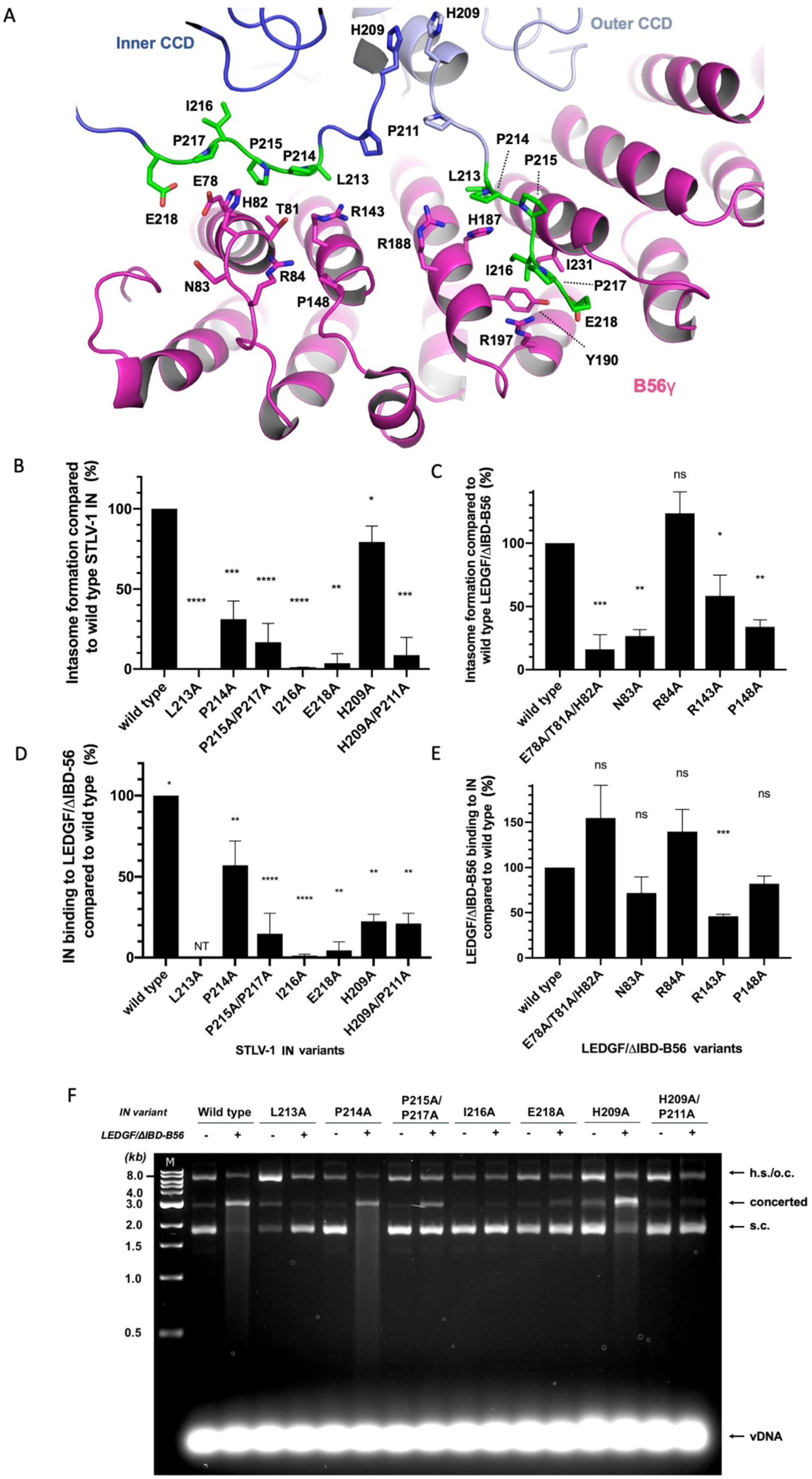
Residues involved in B56γ binding to the STLV-1 intasome. **A**, Two distinct IN binding sites on B56γ are shown binding the conserved SLiM region contained on two IN chains constituting half of the two-fold symmetrical IN dimer of dimers. The inner and outer CCD domains are highlighted in dark and bright blue, respectively, while B56γ is visible in pink. Residues mutated in the following experiments are shown in sticks. A subset of these, belonging to the LxxIxE SLiM motif are coloured in green. **B-C**, The effect of point mutants of STLV-1 IN (**B**) and LEDGF/ΔIBD-B56γ (**C**) on intasome assembly as measured by EMSA (a typical gel is shown in Figure S16). As the presence of LEDGF/ΔIBD-B56γ is crucial for STLV-1 intasome assembly *in vitro*, interface residues important for IN-B56γ binding reduce or abrogate intasome formation. **D-E**, Binding of His_6_-STLV-1 IN point mutants to wild-type (WT) LEDGF/ΔIBD-B56γ, measured by the accumulation of a concerted integration product. Other components of this reaction are indicated as: h.s – half-site integration, o.c. – open circular, s.c. – supercoiled, vDNA – viral donor DNA. Stimulation of catalytic activity is impaired for IN mutants involved in both B56γ interaction interfaces, and the P211A “kink”-inducing mutant – confirming results from EMSA (panel B). Averages and standard deviations of three independent replicates are shown; p-values are as follows: ns (*p* > 0.05), * (*p* < 0.05), ** (*p* < 0.005), *** (*p* < 0.0005), **** (*p* < 0.00005). NT stands for not tested.

Rigid-body docking of the IN and B56γ crystal structures into the cryoEM map provided us with a reliable starting model, which we extended by building the remaining regions *ab initio* (Figure S7B-E). While the canonical IN/CCD dimer observed in crystals could be docked directly into the cryoEM map, the CTDs could only be docked as monomers. We speculate that the dimers and trimers observed in IN/CTD crystals (Figure S10) may be relevant within viral particles prior to vDNA synthesis by reverse transcription. The STLV-1 intasome structure revealed a tetrameric assembly of IN subunits organised around the vDNA ends, with all domains of the tetramer resolved in the cryoEM map (Figure 1). Flanking two sides of the intasome are two B56γ subunits, which resemble epaulettes. The LEDGF parts of the LEDGF/ΔIBD-B56γ constructs are not resolved in the cryoEM reconstruction. Thus, while the LEDGF portion of our B56γ construct helps to chaperon STLV-1 intasome assembly *in vitro*, it is not involved in stable interactions within the resulting nucleoprotein complex.

The tetrameric architecture of the delta-retroviral intasome resembles that of prototype foamy virus (PFV), which is also a tetramer (7) (Figure S12). As was demonstrated by recent structures of lenti- and beta-retroviral intasomes (10,12,13) the oligomeric state of IN within the intasome is dictated by the availability of the CTDs to reach their synaptic positions within the CIC. When the CCD-CTD linker length or topology does not allow for such positioning, the CTDs are provided *in trans* by flanking IN subunits, yielding higher-order IN complexes. The STLV-1 IN CCD-CTD linker length,19 amino acids, is similar to that of lentiviruses (20-22 amino acids) (10) (Table S6). While the lentiviral CCD-CTD linker adopts α-helical conformation (12,27), the corresponding delta-retroviral region is intrinsically disordered (Figure S13). In the cryoEM map, the STLV-1 IN CCD-CTD linker exists in an extended coil conformation providing ample scope for synaptic CTD positioning *in cis* (Figure S14).

B56γ is recruited to the delta-retroviral intasome by the LxxIxE SLiM motif within the IN/CCD-CTD linker (Figure 2A). The LxxIxE consensus sequence is known to provide a binding site for numerous endogenous PP2A interactors and substrates by targeting a conserved groove in the centre of the B56γ subunit of the heterotrimeric PP2A (21,25,26). Delta-retroviruses may have acquired the LxxIxE motif in the course of their evolution to usurp this normal cellular function. Indeed, viruses often employ molecular mimicry to exploit host signalling, and usurping cellular PP2A is not uncommon (28). For some pathogens, such as the Ebola virus, the interaction with PP2A is essential for viral replication (29). Although the LxxIxE-containing region in IN is predicted to be intrinsically disordered, within the structure of the intasome it is stabilised by interactions with B56γ (Figure S13). IN residue Pro211 is highly conserved amongst HTLV/STLV isolates and caps the CCD domain, which introduces a kink in the protein backbone and allows the CCD-CTD linkers in both IN dimers to run perpendicularly to each other providing stable association with B56γ (Figure S13). Unexpectedly, all four SLiM regions in the intasome participate in binding, creating two distinct binding sites for each of the two intasome-recruited B56γ molecules. The previously characterised canonical PP2A SLiM-binding site (26), also resolved in our IN-B56γ crystal structure, involves IN residues Leu213, Ile216 and Glu218 from the outer IN protomer, and B56γ residues His187, Arg188, Tyr190, Arg197, Ile227 and Ile231 (Figure 2A). We previously identified B56γ Arg197 to be critical for binding and stimulating the concerted integration activity of HTLV-1 and HTLV-2 INs (4). In contrast, Arg188, which is important for binding to endogenous phosphorylated substrates (26), is dispensable for binding to delta-retroviral INs (4). Here, we were fascinated to identify an additional novel binding site on B56γ, accommodating the inner IN CCD-CTD linker that runs along the width of B56γ and involves B56γ residues Glu78, Thr81, His82 and Arg143 (Figure 2A). Therefore, the virus uses a remarkable strategy, exploiting the oligomeric assembly of the intasome, to bind to PP2A at two separate sites by means of the same intrinsically disordered yet highly-conserved region, located on neighbouring IN protomers (Figure S13). The presence of a histidine residue, central to the binding at both SLiM-recipient sites, is also of note. Mutations of B56γ residues Glu78, Thr81, His82 and Arg143 to Ala abrogated binding to IN and abolished stimulation of intasome assembly and strand transfer activity (Figure 2B-F). In addition, B56γ residues Asn83 and Pro148, that are in close proximity to the IN CTD, appear to only play a role specific to intasome assembly. Indeed, mutations of these B56γ residues did not affect binding to free IN (Figure 2E), whilst significantly reducing intasome assembly and strand transfer activity (Figure 2C).

It was recently shown that a subset of PP2A-B56 interactors harbour a positively charged motif, complementary to an acidic patch on B56 (30). Delta-retroviral INs do not engage with this acidic patch but appear to have evolved to use an interface employed by other PP2A-B56 interactors. Indeed, Flag-immunoprecipitation of wild type or mutant full-length B56γ from human cells using Ala substitution of Glu78, Thr81, His82, Asn83 and Arg84 showed that these residues are critical for interaction with endogenous PP2A binding partners BUBR1 and CHK2 (21), while not affecting holo-enzyme formation (Figure S15). Of note, not all PP2A-B56 substrates harbour LxxIxE motifs, and some of the SLiM binding partners of PP2A-B56 serve to recruit the phosphatase for dephosphorylation of another macromolecule. For example, BUBR1 recruits PP2A-B56 to kinetochores to dephosphorylate KNL1 and allow mitotic progression (31,32), while the Ebola virus nucleoprotein NP recruits PP2A-B56 to dephosphorylate its viral transcription factor VP30 (29).

HTLV-1 IN forms complexes with the PP2A-B56 holoenzyme (4). The structure reported here contains the regulatory subunit of PP2A and allows modelling of the supramolecular assembly with the entire heterotrimeric phosphatase (Figure 3). Whether phosphatase activity *per se* plays a role in PP2A-B56 modulation of delta-retroviral infection is currently unknown. However, provided that the missing IN NTD-CCD linker of the outer IN protomer does not preclude access to the active site, phosphorylated peptides could still be substrates of this supramolecular assembly.

**Figure 3.**
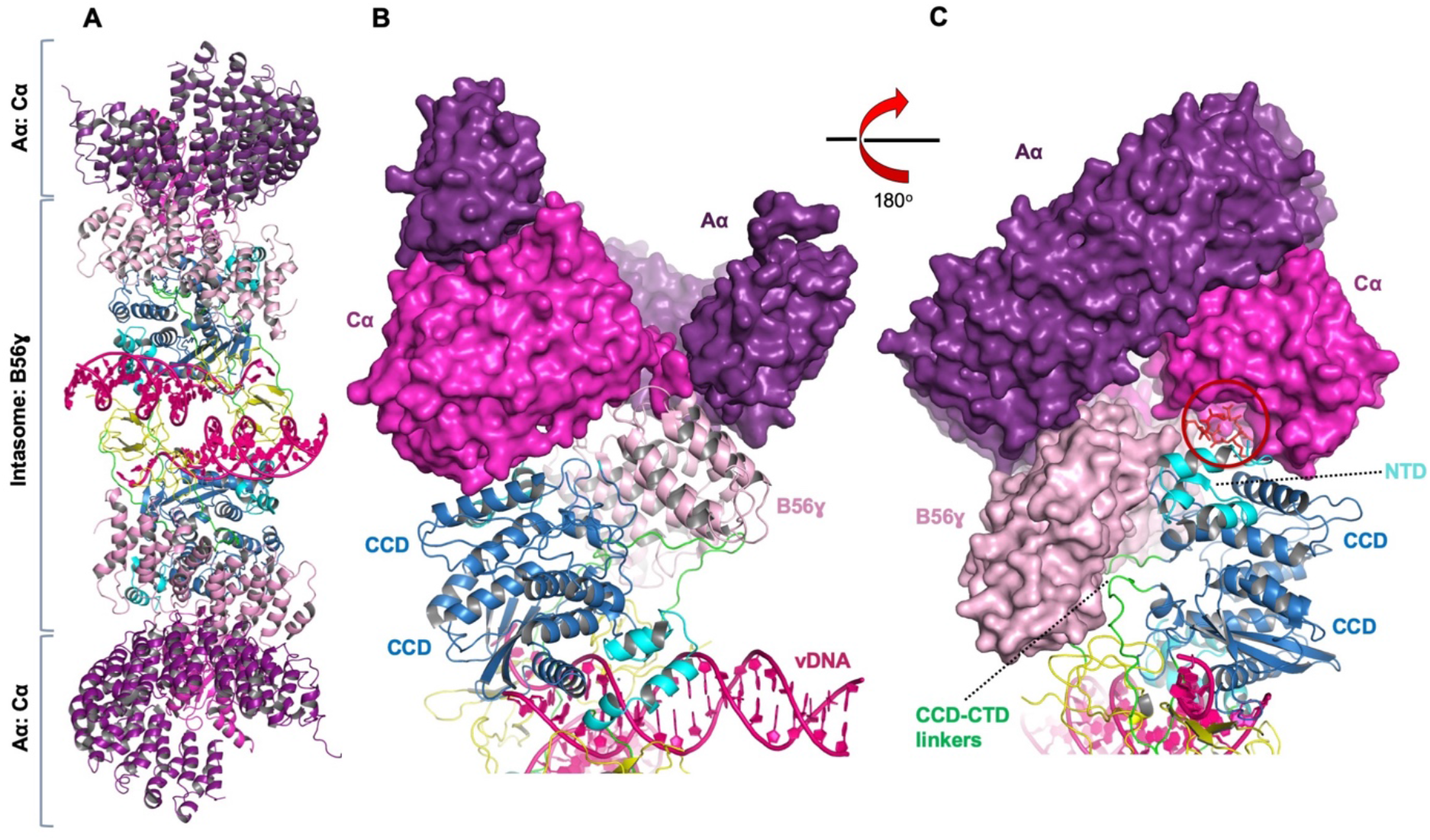
Model of the complete STLV-1 intasome: PP2A holoenzyme complex. **A**, The model was created by conformational alignment of the crystal structure of PP2A holoenzyme (PDB ID 2IAE) and the cryoEM structure of the STLV-1 intasome: B56γ complex using the B56γ subunit as a common component of both structures. **B**, After rigid-body modelling, no clashes are seen between any atoms of the cryoEM intasome structure and the PP2A holoenzyme. Interactions between other subunits of PP2A and IN may also take place in the context of the intasome: PP2A complex. **C,** A molecule of microcystin, a peptide mimetic, co-crystallised in the PP2A holoenzyme structure is illustrated in red sticks. The modelling suggests that in the context of the STLV-1 intasome: PP2A holoenzyme complex the phosphatase active site (circled in red) could remain open for substrate binding.

Recruitment of the holoenzyme greatly expands the surface of the nucleoprotein complex and is expected to provide the delta-retroviral integration machinery an interface for chromatin-bound PP2A interaction partners (21–24). HTLV-1 gains access to chromatin during mitosis, when the bulk of chromatin is highly condensed. Thus, interaction with PP2A may allow the virus to locate chromatin loci that are bookmarked for expression soon after completion of cell division (33). Our structural work, presented here, will be instrumental to fully characterize the role of PP2A-B56 in HTLV-1 infection, and given that small changes in the active site between different retroviral INs significantly impact drug binding (34), forms the foundation to develop highly specific inhibitors of HTLV-1 integration.

**Supplementary Information** is linked to the online version of the paper at www.nature.com/nature.

## Acknowledgements

We thank Dr R. Carzaniga and L. Collinson for the maintenance of Vitrobot and Tecnai G2 microscope and user training; Drs A. Purkiss, P. Walker and M. Oliveira for computer and software support; Drs C. McAuley, P. Romano, D. Hall and J. Beale for their help and assistance at the Diamond Lightsource Synchrotron, Dr M. Morgan (Imperial College London) for expert help with in-house crystallisation screening and data collection, and N. Cook (Francis Crick Institute) for expert help with cryoEM grid vitrification. We are grateful to Dr. A. Engelman (Dana-Farber Cancer Institute) for helpful comments and critical reading of the manuscript and Dr. A. L. B. Ambrosio (Laboratório Nacional de Biociências, Brazil) for the generous gift of pET28a-SUMO. This work is supported by the Wellcome Trust (Investigator Award to G.N.M., 107005/Z/15Z) and the Royal Society (RG120032, to G.N.M.). Work in P.C. laboratory is supported by the Francis Crick Institute, which receives its core funding from Cancer Research UK (FC001061), the UK Medical Research Council (FC001061), and the Wellcome Trust (FC001061).This article is independent research funded by the National Institute for Health Research (NIHR) Imperial Biomedical Research Centre (BRC). The views expressed in this publication are those of the authors and not necessarily those of the NHS, the National Institute for Health Research or the Department of Health.

## Author Contributions

G.N.M. discovered how to assemble STLV-1 intasomes, assembled, purified and prepared samples for negative-stain EM; G.N.M. and P.C. collected and analysed negative stain EM data; G.N.M. conducted the Flag-IP experiments; M.S.B. purified all wild type and mutant full-length proteins, assembled intasomes for biochemical analysis and cryo-EM data collection; M.S.B. purified, crystallised, solved and refined the X-ray structures of the individual IN domains; M.S.B. and P.C. analysed the cryoEM data; M.S.B., P.C. and V.E.P refined the atomistic model; J.J.M. optimized the assembly of the IN(200-297):B56γ(11-380) complex, conducted crystallization screening, optimization, X-ray data collection; M.S.B, J.J.M. and G.N.M refined the structure of the IN: B56γ complex; Z.H. prepared graphene oxide carbon and prepared the grids for cryoEM data collection; A.N. collected the cryoEM data; M.S.B, J.J.M., P.C. and G.N.M prepared the manuscript with contributions of all authors.

## Author information

The crystal structures have been deposited with the Protein Data Bank and are available under the following identifiers: HTLV-2/CCD-Mg^2+^ (dimeric form): 6QBV; HTLV-2/CCD-Mg^2+^ (trimeric form): 6QBT; HTLV-2/CCD-Ca^2+^ (dimeric form): 6QBW; HTLV-1/CTD: 6TJU; HTLV-1 IN(200-297)-B56γ: 6TOQ. Raw diffraction images are available upon request. The cryoEM structure has been deposited with the Protein Data Bank and EMDB and are available under the following identifiers 6Z2Y and EMD-11052. Reprints and permissions information is available at www.nature.com/reprints. The authors declare no competing financial interests. Correspondence and requests for materials should be addressed to G.N.M.

The authors declare no competing interests.

