## Supplementary Information for "CRYO-EM STRUCTURE OF THE DELTA-RETROVIRAL INTASOME IN COMPLEX WITH THE PP2A REGULATORY SUBUNIT B56γ"

**Supplementary Information for manuscript "Cryo-EM structure of the delta-retroviral intasome in complex with the PP2A regulatory subunit B56γ." by Barski M et al.**

### Materials and Methods

#### Cloning of HTLV-1, HTLV-2, STLV-1 IN and human PPP2R5C constructs for prokaryotic expression

The pET28a(+)-SUMO-H6P vector was engineered by ligating annealed oligonucleotides JM1 and JM2 into *Bam*HI/*Eco*RI-digested pET28a(+)-SUMO plasmid (a gift of A. L. B. Ambrosio, Laboratório Nacional de Biociências, Campinas, Brazil). DNA sequence corresponding to residues 53 to 221 of HTLV-2 IN was PCR-amplified using primers GM130 and MB094. Following *Mfe*I and *Sal*I digest, the amplicon was ligated into EcoRI/SalI-digested pET28a(+)-SUMO-H6P. pET28(+)-SUMO-H6P-HTLV-1 IN(228-286) and (200-297) were generated by PCR amplification of the corresponding coding sequence using primers MB114 and MB054 and GNM378 and GM110, respectively. *Eco*RI- and *Sal*I-digested amplicons were ligated into similarly digested plasmid DNA. The STLV-1 MarB43 IN coding sequence was generated by codon optimizing the *integrase* gene (corresponding to nucleotides 4338-5231 of the STLV-1 MarB43 isolate genome, GenBank ID: AY590142.1) for eukaryotic expression. The codon optimized DNA was PCR-amplified using primers GNM747 and GNM748, digested with *Eco*RI and *Xho*I and ligated into similarly digested pET28a(+)-SUMO-H6P. pET28a(+)-SUMO-PPP2R5C(11-380) has been previously reported (1). To express $His_6$-SUMO-LEDGF/ΔIBD-B56γ(11-380) the bases corresponding to amino acids 1-324 of *PSIP1* (gene encoding LEDGF) were amplified using primers GNM641 and GNM642, digested with *Bam*HI and *Eco*RI and ligated into *Bam*HI/*Eco*RI-digested pET28(+)-SUMO-PPP2R5C(11-380). All primers and their sequences are listed in Table S1; all DNA constructs reported in this work were verified by sequencing.

#### STLV-1 IN and LEDGF/ΔIBD-B56γ site-directed mutagenesis

Two-step splicing PCR was used to generate site-directed mutants of STLV-1 IN full-length, and LEDGF/ΔIBD-B56γ(11-380). The following primer pairs were used for cloning SLTV-1 IN mutants L213A (primer GNM771, GNM770 and GNM769, GNM772), P214A (GNM773, GNM770 and GNM774, GNM769), P215A/P217A (GNM775, GNM770 and GNM776,

GNM769), I216A (GNM777, GNM770 and GNM778, GNM769), E218A (GNM779, GNM770 and GNM780, GNM769), H209A (GNM769, GNM824 and GNM770, GNM829), H209/P211A (GNM769, GNM826 and GNM770, GNM830). The PCR products were then spliced together using primers GNM770 and GNM769, digested with *Eco*RI/*Xho*I and ligated into similarly digested pET28(+)-SUMO-H6P. The following primer pairs were used to clone LEDGF/ΔIBD-B56γ mutants E78A/T81A/H82A (GNM283, GNM641 and GM142, GNM750), R143A (GNM283, GNM751 and GM142, GNM752), N83A (GNM283, GNM844 and GM142, GNM845), R84A (GNM283, GNM833 and GM142, GNM834), P148A (GNM283, GNM837 and GM142, GNM838). The PCR products were then spliced together using primer pairs GM142 and GNM283, digested with *Eco*RI/*Sal*I and ligated into *Eco*RI/*Sal*I-digested pET28(+)SUMO-LEDGF/ΔIBD-PPP2R5C(11-380).

**Cloning of full length *PPP2R5C* for eukaryotic expression.**

The full length *PPP2R5C* gene was PCR-amplified from HeLa cDNA using primers GNM380 and GNM367, digested with *Age*I and *Sal*I and ligated into *Age*I/*Xho*I-digested pQFlag puroR (2). Sequencing of the resulting construct established that cloned cDNA corresponds to isoform 2 of *PPP2R5C* gene transcript. Point mutations were introduced by splicing PCR as described above with the difference that the primers annealing to the 5' and 3' ends respectively were JM3 and GNM380.

**Crystallisation of HTLV-2 IN/CCD (53-221)**

Expression was conducted in *E. coli* Rosetta-2 (DE3) pLacI cells (Novagen) in Terrific Broth (TB, Melford). Cells were grown to the $OD_{600}$ of 2.0 at 30°C, followed by 30-min incubation at 18°C and induction by 0.01% IPTG at 18°C overnight. The pelleted cells were disrupted by sonication in 0.5 M NaCl, 10 mM imidazole, 50 mM Tris-HCl pH 7.4, 1 mM phenylmethylsulfonyl fluoride (PMSF). The soluble protein, captured on His-Select nickel immobilized metal affinity chromatography (IMAC) resin (Sigma-Aldrich, UK), was extensively washed with buffer containing 10 mM imidazole, 0.5 M NaCl, 50 mM Tris-HCl pH 7.4. Following elution from IMAC with the wash buffer supplemented with 200 mM imidazole, cleavage was conducted at 4°C overnight with HRV 3C protease (using 1 mg protease per 20 mg protein) in presence of 5 mM DTT. Subsequently, the protein was diluted 5-fold with salt-free buffer to achieve a final NaCl concentration of 100 mM and subjected to cation-exchange chromatography on a 5-ml high-performance SP column (GE Healthcare,

UK). Fractions containing IN/CCD were pooled and injected onto an S200 16/60 size-exclusion column (GE Healthcare, UK) with 50 mM Tris-HCl pH 7.4, 0.5 M NaCl, 2 mM DTT as running buffer. Positive fractions were collected and concentrated in a 10-kDa MWCO Vivaspin centrifugal ultrafiltration device (Sartorius) to a volume of 1 mL. This was then diluted two-fold in ice-cold ultra-pure water and concentrated further to yield 15 mg/mL HTLV-2 IN/CCD in 250 mM NaCl, 1 mM DTT, 25 mM Tris-HCl pH 7.4.

The sample prepared as above was used to set up 960 sitting-drop vapour diffusion sparse-matrix crystallisation conditions. Initial screens were set up using the Mosquito liquid dispenser (SPT Labtech), with 800 nL drops consisting of a 1:1 ratio of protein-to-precipitant and allowed to incubate at 18°C. Final crystal growth was achieved by hanging-drop vapour diffusion in 2 μL drops consisting of 2:1 ratio of 21 mg/mL HTLV-2 IN/CCD-to-precipitant. For the CCD-$Mg^{2+}$ complex, the precipitant solution comprised of 100 mM Tris-HCl pH 8.5, 14% polyethylene glycol (PEG) 8,000 and between 100 mM and 250 mM $MgCl_2$. For the CCD-$Ca^{2+}$ complex, the precipitant used was: 100 mM Tris-HCl pH 8, 18% PEG 6K, 250 mM $CaCl_2$. Maximum crystal growth in the condition containing $MgCl_2$ and $CaCl_2$ appeared after 48 h and 3 months at 18°C incubation, respectively. Crystals grew as large elongated hexagons or cubes, with the lengths of up to 0.5 mm.

Crystals, cryoprotected in a step-wise fashion with reservoir solution supplemented with 10%, then 20% glycerol (v/v), were cryo-cooled by plunging into liquid nitrogen. Native data collection was carried out at beamline I03 for the IN/CCD-$Mg^{2+}$ and I04 for the IN/CCD-$Ca^{2+}$ at Diamond Light Source (Didcot, UK). A 0.976 Å wavelength beam was used to collect diffraction images with 0.01 s exposures under 0.1° oscillation angle, over a total rotation angle of 360°. Crystals diffracted to a maximum resolution of approximately 2 Å. Data were indexed and integrated in Xia2 (3) using XDS (4), which identified datasets belonging to two unique space groups. Datasets were scaled and merged in CCP4i2 (5) Aimless (6), with the resolution limit adjusted accordingly until satisfactory signal-to-noise and completeness were reached. Unit cell composition was estimated from Matthew's coefficient. Phases were obtained by molecular replacement in PHASER (7) through the PHENIX software suite (8). The mouse mammary tumour virus (MMTV) IN/CCD (PDB ID: 5CZ1) was used as a search model and resulted in a solution with a TFZ score of 50. PHENIX Autobuild (9) was successful in building the majority of residues in all chains. The remaining residues were added manually in Coot and the model was refined in Refmac 5.8.

**Crystallisation of HTLV-1 IN/CTD (228-286)**

The HTLV-1 IN/CTD construct that led to good quality highly diffracting crystals encodes for residues 228-286 and is further referred to as the IN/CTD. Expression was conducted as described above for HTLV-2 IN/CCD. Extraction was performed in 50 mM Tris-HCl pH 7.4, 0.5 M NaCl, 10 mM imidazole, 1 mM PMSF, through sonication of the resuspended. This was then clarified by centrifugation at 50 000 *g* for 15 min at 4ºC. Once bound to the IMAC resin, the sample was washed with 100 mL of 50 mM Tris-HCl pH 7.4, 2 M NaCl, 10 mM imidazole in order to dissociate the contaminating nucleic acid. Following elution in 50 mM Tris-HCl pH 7.4, 0.5 M NaCl, 200 mM imidazole, digestion of the $His_6$-SUMO-$His_6$ tag was performed with HRV 3C protease overnight at 4ºC in presence of 5 mM DTT. Due to the apparent cold-induced precipitation of HTLV-1 IN/CTD, all the following steps were performed at room temperature. Ion-exchange chromatography was conducted with a high-performance SP column (GE Healthcare, UK) with the sample diluted to 150 mM NaCl concentration for binding. Gel filtration of positive fractions was carried out on Superdex-200 16/60 size-exclusion column (GE Healthcare, UK) with 50 mM Tris-HCl pH 7.4, 300 mM NaCl. Positive fractions were pooled, diluted 1:1 with ultrapure water to yield final buffering conditions of 25 mM Tris-HCl pH 7.4, 150 mM NaCl, and supplemented with 2 mM DTT. The sample was concentrated in a 3K MWCO Vivaspin ultrafiltration device (Sartorius) to 14 mg/mL.

A Mosquito liquid dispenser was used to set up 960 crystallisation conditions in sitting-drop plates. Successful crystallisation was observed in several conditions, of which the most promising one comprised 1.3 M ammonium tartrate dibasic, 0.1 M bis-tris propane (BTP)-HCl pH 7.0. Rhombohedral crystals appeared after three days. Optimisation led to the selection of 1.1 M ammonium tartrate dibasic as the optimal condition for crystal growth in the hanging-drop format, and the protein:precipitant ratio of 2:1. Apparent over-nucleation, affecting crystal size, was resolved by microseeding. Seeds were prepared by maceration of medium-size crystals in mother liquor containing the same crystallisation components as the target condition. Seeds were then collected, diluted 1,000-fold in the reservoir solution and kept on ice. Streak-seeding was performed immediately after.

Crystals were harvested and cryoprotected in a step-wise fashion with reservoir solution supplemented with 10%, then 20% glycerol (v/v). Crystals were then cryo-cooled in liquid nitrogen. Native data collection was carried out at beamline I04 at Diamond Light Source (Didcot, UK). A 0.979 Å wavelength beam was used to collect diffraction images with 0.01 s exposures per 0.1° oscillation angle, over a total rotation angle of 360°. Crystals diffracted to a maximum resolution of 1.14 Å. Data were indexed and integrated in XDS (4) via Xia2 (3). Datasets were scaled and merged in CCP4i2 (5) using Aimless (6), with the resolution limit adjusted accordingly until satisfactory signal-to-noise and completeness were reached. Unit cell composition was estimated from Matthew's coefficient. Phases were obtained by molecular replacement using PHASER (7) through the PHENIX software suite (8). Using the MVV IN/CTD structure as a replacement model (PDB code 5LLJ) resulted in a single solution with a TFZ score of 30.1. PHENIX Autobuild (9) was successful in building the majority of residues in all chains. The remaining residues were added manually in Coot and the model was refined in Refmac 5.8.

**Crystallisation of HTLV-1 IN (200-297) : B56γ complex**

The PP2A regulatory subunit B56γ (11-380) (further referred to as B56γ) was expressed and purified as described previously (10). Expression and purification of HTLV-1 IN (200-297) was performed as described above for the 228-286 construct. HTLV-1 IN (200-297) did not display cold-induced precipitation properties and was therefore kept cold throughout the purification.

Freshly purified B56γ and HTLV1 IN (200-297) were concentrated to 5 mg/mL, and combined at a 1:1 molar ratio, followed by incubation on ice for 10 minutes. Samples were dialysed against ice-cold buffer (50 mM Tris-HCl pH 8.0, 200 mM NaCl, 2 mM DTT) overnight at 4˚C. The complex was purified by gel filtration in the above buffer and the peak fractions were confirmed to contain both proteins by SDS-PAGE. The HTLV-1 IN (200-297)-B56γ complex was concentrated to 26 mg/mL in a 10 kDa MWCO ultrafiltration device (Vivaspin). A 100 nL sample and 100 nL crystallisation buffer drops were set up in a sitting-drop vapour diffusion format using the mosquito liquid-handling robot. Initial crystallisation was observed in 0.1 M Na/$KPO_4$ pH 6.2, 25% v/v 1,2-propanediol, 10% v/v glycerol. Optimisation of this condition led to crystallisation in the hanging-drop vapour-diffusion format in 0.1 M Na/KPO4 pH 6.2, 20% 1,2-propanediol, 10% glycerol. In order to increase crystal size and control nucleation, microseeding was conducted with crystals from the above condition. Crystals suspended in mother liquor were macerated with the crystal crusher tool (Hampton Research), and a 1:100 dilution of the seed stock was streak-seeded onto new drops. The resulting crystals were thin plates, approximately 50 μm by 50 μm.

Data were collected on beamline I24 at Diamond Light Source (Didcot, UK). Reflections were indexed using Xia2 (3) in 3dii mode (4). In order to increase the signal-to-noise ratio, two wedges of images with the highest Bragg reflection number were extracted from the original dataset and re-indexed together in space group $P4_32_12$. Phaser (7) was used for phasing by molecular replacement using the B56γ structure (PDB ID: 2JAK) as the template. PHENIX Autobuild (9) was successful in building the majority of residues in all chains. The remaining residues were added manually in Coot and the final model was refined in Refmac 5.8.

**Expression and purification of full-length STLV-1 IN**

Expression was conducted as described above for HTLV-1 IN constructs. Next, cells were resuspended in 25 mM Tris-HCl pH 7.4, 1 M NaCl, 7.5 mM CHAPS, 1 mM PMSF, 10 mM imidazole, and 20 μg/mL lysozyme. Cells were then lysed by sonication and clarified by centrifugation at 50, 000 *g* for 15 min at 4°C. Following Ni-assisted IMAC, conducted as for HTLV-1 IN constructs, cleavage of the SUMO solubility tag was conducted with either HRV 3C protease for the non-$His_6$-tagged product or Ulp-1 protease for the $His_6$-tagged product at 4°C overnight, in presence of 5 mM DTT. IN concentration was kept below 2 mg/mL to prevent aggregation and precipitation. Ion-exchange chromatography was then conducted on a high-performance SP column (GE Healthcare, UK) following binding of the sample diluted with buffer without NaCl to achieve a final NaCl concentration of 250 mM NaCl. Peak fractions, eluted with a NaCl gradient, containing pure IN were then pooled and injected onto a Superdex 16/60 size-exclusion column (GE Healthcare), pre-equilibrated in 25 mM Tris-HCl pH 7.4, 7.5 mM CHAPS and 1 M NaCl. Fractions containing pure IN were then pooled and dialysed in a 10K MWCO Snakeskin dialysis tubing (Life Technologies) against 20 mM BTP-HCl pH 6, 1 M NaCl, 2 mM DTT, at 4°C overnight. Following completion of dialysis, the sample was recovered and concentrated in a 10K MWCO ultrafiltration device (Vivaspin) to a concentration of 2 mg/mL or higher. For storage, glycerol was added to a final concentration of 10%, the sample was flash frozen in liquid nitrogen and stored at -80°C until needed. Note, the $His_6$-tagged IN proteins were less soluble than the untagged versions of the protein and yields of $His_6$-IN(L213A) were too low for use in binding assays.

**Expression and purification of LEDGF/ΔIBD-B56γ**

LOBSTR RIL cells (11) (Kerafast) were used for expression of LEDGF/ΔIBD-B56γ. Cells were grown in LB to an $OD_{600}$ of 0.6 and following induction with 0.01% IPTG were further incubated at 25°C for 3 hours. The temperature was then lowered to 16°C and induction was continued overnight. Cells were resuspended in a solution containing 50 mM Tris-HCl pH 8, 1 M NaCl, 10 mM imidazole, 20 μg/mL lysozyme and 1 mM PMSF. Following sonication to disrupt cells, the extract was clarified by centrifugation. IMAC was performed on a Ni-NTA column (GE Healthcare, UK). Thorough wash was performed in 50 mM Tris-HCl pH 8, 1 M NaCl, 10 mM imidazole. The last wash was performed in 50 mM Tris-HCl pH 8, 0.5 M NaCl, 10 mM imidazole, followed by elution with a buffer containing 25 mM Tris-HCl pH 8, 0.5 M NaCl, 200 mM imidazole. Cleavage was performed overnight with Ulp1 SUMO protease in presence of 5 mM DTT. The protein was diluted with salt-free buffer to achieve NaCl concentration of 125 mM, injected into an HP Q column (GE Healthcare, UK) and eluted with a gradient of 0.15 - 0.5 M NaCl. Fractions containing LEDGF/ΔIBD-B56γ were pooled and the protein was polished by size exclusion chromatography through a Superdex S200 16/60 gel-filtration column (GE Healthcare, UK) operated in 300 mM NaCl, 25 mM Tris-HCl pH 8. Fractions containing pure LEDGF/ΔIBD-B56γ were supplemented with 2 mM DTT and concentrated to 20 mg/mL using a 30-KDa MWCO ultrafiltration device (Vivaspin). Protein, supplemented with 10% glycerol, was flash-frozen in liquid nitrogen and stored at -80°C until further use.

**STLV-1 strand-transfer activity assays**

Assays were conducted using purified recombinant STLV-1 IN and the vDNA LTR oligonucleotide mimics (Table S2) were annealed in 100 mM Tris-HCl pH 7.4, 400 mM NaCl. The optimised reaction conditions were: 25 mM BTP-HCl pH 6, 2.5 μM STLV-1 IN, 2 μM vDNA, 60 mM NaCl, 13.28 mM DTT, 10 mM $MgCl_2$, 10 μM $ZnCl_2$. After addition of vDNA, the reaction was incubated at 37°C for 10 min. Following a co-incubation with supercoiled target DNA (s.c. tDNA) for one hour at 37°C, samples were processed as described previously (12). Where IN : LEDGF/ΔIBD-B56γ was used, IN was pre-incubated with 1:2 ratio of LEDGF/ΔIBD-B56γ to STLV-1 IN for 30 min at 4°C prior to incubation with vDNA.

**Electrophoretic mobility shift assays (EMSA)**

Five μL STLV-1 IN (1.6 mg/mL in 20 mM BTP-HCl pH 6, 1 M NaCl, 2 mM DTT) was mixed with 5 μL 3.84 mg/mL IN : LEDGF/ΔIBD-B56γ and diluted with 20 μL buffer to yield a final NaCl concentration of 200 mM. Following incubation at 4°C for 30 min, the samples were supplemented with 0.5 μL 20 μM Atto680-labeled vDNA (Table S2), and the reaction volume increased to yield a final NaCl concentration of 60 mM. The reaction was placed at 37°C for 10 min, then NaCl concentration was increased to 1.2 M and allowed to equilibrate at room temperature. Samples, supplemented with 10 μg/mL heparin, were separated on a 3% low melting point agarose gel containing 10 μg/mL heparin (13). Densitometry of bands corresponding for the intasome was carried out in ImageJ. Measurements from at least three independent experiments were taken, standard deviations and *p*-values were calculated in Prism 8.

**Pull-down assays**

$His_6$-tag pull-down assays were done as previously described (1). Densitometry of bands was carried out in ImageJ. Measurements from at least three independent experiments were taken. Values for the pulled-down IN or LEDGF/ΔIBD-B56γ in each condition were normalised to the quantity of the bait present. Standard deviations and *p*-values were calculated in Prism 8.

**Assembly and purification of the STLV-1 intasome : LEDGF/ΔIBD-B56γ nucleoprotein complex**

The STLV-1 IN : LEDGF/ΔIBD-B56γ complex was first assembled by mixing equimolar (0.03 mmol) quantities of IN and LEDGF/ΔIBD-B56γ and dialysing overnight at 4°C against 0.5 L of 25 mM Tris-HCl pH 7.4, 200 mM NaCl, 2 mM DTT. We have previously found (see section Crystallisation of HTLV-1 IN (200-297) : B56γ) that this condition promotes IN : LEDGF/ΔIBD-B56γ complex formation. This sample was then concentrated at 4°C, 1,935 *g* in a 30-KDa MWCO ultrafiltration device (Vivaspin) to a concentration of 0.2 mM. A mixture containing 0.7 mL 20 mM BTP-HCl pH 6, 10 mM $CaCl_2$, 10 mM DTT, 10 μM $ZnCl_2$ and 25 μM STLV-1 U5 S30 double-stranded vDNA (Table S2) was placed in a heat block set to 37°C and incubated for 10 min. The previously prepared IN : LEDGF/ΔIBD-B56γ complex was then added, mixed by gently flicking the tube and the tube placed back in the heat block for 10 min. Upon addition of the protein complex and during the course of incubation dense, white precipitate appeared. Following the incubation, the precipitate was dissolved by addition of NaCl to a final concentration of 1.2 M, gentle up-and-down mixing, and a further 15 min incubation at room temperature. Increasing NaCl concentration allowed for complete dissolution of the precipitate and recovery of assembled nucleoprotein complex. The sample was then immediately loaded onto an S200 10/300 Increase size-exclusion column (GE Healthcare, UK). For samples prepared for negative-stain observations, the size-exclusion mobile phase was 20 mM BTP-HCl pH 6, 1.2 M NaCl. For cryoEM preparations, the size-exclusion mobile phase was 20 mM BTP-HCl pH 6, 0.3 M NaCl. Peak fractions were pooled and tested for integration activity in the presence of 10 mM $MgCl_2$ and 300 ng target DNA (supercoiled pGEM-9Zf(−)), as well as by SDS-PAGE. Fractions corresponding to the highest strand transfer activity were used for negative-stain and cryoEM grid preparation.

**Negative stain grid preparation, data collection and processing**

Four µl drops of freshly assembled and purified STLV-1 intasomes were spotted on carbon-coated 300-mesh copper grids (EM Resolutions, catalogue #C300Cu), which had been glow-discharged for 30 s at 45 mA using an Emitech K100X instrument (EMS) and allowed to bind for 1 min. Excess sample was blotted and absorbed particles were stained with 2% uranyl acetate. Grids were imaged on a Tecnai G2 Spirit LaB6 transmission 120-kV electron microscope (Thermo Fisher Scientific) with an Ultrascan-1000 camera (Gatan) at 30,000x magnification, resulting in a magnified pixel size of 3.45 Å. A total of 152 micrographs were taken, from which 22,000 particles were picked using EMAN2. 2D classification was done in Relion-2 and 8,790 particles were used for *ab initio* 3D reconstruction and homogenous refinement.

**CryoEM grid preparation and data collection**

Four µl freshly prepared intasome ($A_{260}$ ~1.5, corresponding to ~ 2.3 µM nucleoprotein complex) was applied on C-flat holey carbon gold grids (Electron Microscopy Sciences, catalogue #CF-1.2/1.3-4Au) or on UltraAuFoil R 1.2/1.3 grids (14) (Electron Microscopy Sciences, catalogue Q350AR13A) that were freshly-coated with graphene oxide following a published protocol (15). Glow-discharging was avoided in both cases. The use of fresh C-flat grids was essential to observe intasomes in open holes. The grids, incubated for 1 min at 22°C and 95% humidity, were blotted for 2-3 s prior to plunge-freezing in liquid ethane using a VitroBot Mark IV instrument (Thermo Fisher Scientific). Data were collected on Titan Krios electron microscope operating at 300 kV with a Falcon III direct electron detector in counting mode (Thermo Fisher Scientific). A pixel size of 1.09 Å and defocus range of -1.6 to -3.6 μm was used for the data collections. A total electron exposure of 34 e/Å$^2$ was fractionated across 30 movie frames over a 60 second exposure time. A total of 8,088 and 8,949 movies were recorded from open hole C-flat (OH dataset) and graphene oxide supported UltrAuFoil (GO dataset) grids, respectively with EPU 1.9.0 software (Thermo Fisher Scientific).

**Single-particle image processing and 3D reconstruction**

Micrograph movie frames were aligned and summed with dose weighting applied as implemented in MotionCor2 (16), and the contrast transfer function (CTF) parameters were estimated from the frame sums using Gctf-v1.06 (17). Following removal of images with evidence of crystalline ice contamination and/or those lacking graphene oxide, 8,022 (OH dataset) and 8,049 (GO dataset) aligned micrographs were retained for particle picking and further image processing. A small subset of micrographs were picked manually with EMAN2 boxer and subjected to reference-free classification in Relion-2 to generate initial 2D class averages in (Figure S4). These were used as templates for picking the entire datasets with Gautomatch v0.56 (http://www.mrc-lmb.cam.ac.uk/kzhang/), resulting in the initial subsets of 2,198,454 (OH dataset) and 2,157,654 (GO dataset) particles. The particles extracted in Relion-3.0, binned by a factor of 2, were subjected to two rounds of reference free 2D classification in CryoSPARC-2. Particles belonging to well-defined 2D classes (599,700 and 493,665 particles for OH and GO datasets, respectively) were subjected to 45 cycles of 3D classification into 17 (OH) or 13 (GO) classes in Relion-3.0 without imposing symmetry, with an initial model generated in CryoSPARC-2. The procedure yielded a single high-resolution class from each dataset. Particles belonging to the best 3D classes (94,517 and 67,397 from OH and GO dataset, respectively) were re-extracted as full-sized images. 3D reconstructions generated from the individual datasets resulted in highly anisotropic maps due to severe preferential orientations of the single particles (Figure S6). Since 3D-FSC analysis (18) indicated favourable complementarity of the data (Figure S6), the datasets were merged and refined as separate optics groups in Relion-3.1. 3D reconstruction, followed by Bayesian polishing, per-particle defocus and beam tilt refinement, as implemented in Relion-3.1, resulted in the final map with minimal anisotropy. 3D-FSC sphericity index of the final map was 0.967 (Figure S6). Gold-standard Fourier shell correlation (FSC) = 0.143 criterion (19,20) was used to estimate resolutions of the 3D reconstructions (Table S5). Local resolution of the cryoEM map was estimated using Blocres from the Bsoft software package (21).

**Integrative model building and refinement**

The quality of the cryoEM map was marginally improved using Resolve density modification procedure (22) implemented in Phenix 1.18-3845, which increased estimated resolution of the reconstruction by 0.15 Å. Density modification was performed under default parameters, using half-maps and macromolecular sequence as inputs. Alternatively, cryoEM map was sharpened using a global B factor (-143 $Å^{-2}$, determined automatically) or locally filtered using post-processing procedure implemented in Relion-3.1 Initially, X-ray crystal structures were docked into resulting cryoEM maps as rigid bodies in Chimera (23). A high-quality homology model of the STLV-1 IN/NTD was generated by SWISS-MODEL server (24). *Ab initio* building residues not present in docked models but resolved in the cryoEM density and manual refitting of docked models was conducted in Coot (25). The globally and locally sharpened maps and density modified map were used to guide model building. The initial model, comprising chains A, B, C, K, and L, was subjected to molecular dynamics structural fitting using Namdinator (26). This model was further adjusted in Coot before the model was duplicated to form chains D, E, F, M, and N which were docked in place using Chimera and rigid body fitted in Coot. The model was manually checked again for clashes between the NCS chains before final real space refinement using Phenix version 1.18-3845 and the density modified map, implementing secondary structure and base pair/base stacking definitions based on the model, metal bond restraints and NCS constraints for the two halves of the symmetric nucleoprotein assembly. Quality of the final atomistic model was assessed with MolProbity (27) and EMRinger (28) (Table S5).

**Tissue culture and Flag-immunoprecipitation (IP)**

Human Embryonic Kidney SV40 large T cells (further referred to as 293T) were cultured in Dulbecco's Modified Eagle Medium (Sigma) supplemented with 10% foetal bovine serum (Sigma), 100 IU/mL penicillin, and 100 μg/mL streptomycin (Sigma) and grown in a humidified incubator at 37°C with 5% $CO_2$. 293T cells were transfected with 20 μg plasmid DNA by calcium phosphate precipitation as previously described (29). Thirty-six hours post-transfection, cells were harvested and moved to ice. All further procedures were done on ice or at 4°C. Cells were lysed in 5 volumes of PP2A IP buffer (20 mM Tris-HCl pH 8.0, 0.1% (v/v) Nonidet P-40, 150 mM NaCl, 3 mM EDTA, 3 mM EGTA) supplemented with Complete EDTA-free protease inhibitor cocktail (Roche), left on ice for 10 min and cellular debris were removed by centrifugation at 16,000 *g* for 30 min. Protein concentration in the extract was determined using the DeNovix spectrophotometer (assuming 1 $A_{280}$ corresponds to 1 mg/ml protein) and 3 mg of total protein for each sample in a volume of 400 μl was allowed to bind to 25 μl pre-washed anti-Flag agarose beads (Sigma) by end-over-end rocking at 4°C. Beads were washed 4 times in 1 ml IP buffer. After removing all remaining liquid from the beads, proteins were eluted by boiling in 30 μl of Laemmli buffer. To detect BUBR1 and CHK2 in the protein lysate, 30 μg total protein (equivalent with 1% used for the IP) was loaded on gel. For detection of Flag-B56γ, Aα and Cα, 10 μg of lysate was loaded. Ten μl eluate was separated on an 11% SDS-PAGE denaturing gel, proteins were electrotransferred onto nitrocellulose. Membranes were blocked in 5% milk/PBS and probed with the following antibodies: horse radish peroxidase (HRP) conjugated mouse anti-Flag antibody (1: 2,000, clone M2, Sigma, A8592), rabbit anti-BUBR1 (1:1,000, Bethyl Laboratories, A300-365A-T), rabbit anti-CHK2 (1:200, Santa Cruz Biotech, H-300, sc-9064), rat anti-Aα (1:2,000 clone 6G3, Insight Biotechnology, sc-56954), mouse anti-Cα (1:2,000, clone 46, Becton-Dickinson, 610556). All antibodies were diluted in 5% milk/PBST (PBS supplemented with 0.1% Tween-20). HRP conjugated donkey anti-rabbit (GE Life Sciences, GE NA934), rabbit anti-rat (Abcam, ab6734) and rat anti-mouse (Abcam, ab131368) were used at 1:2,000 dilution. Detection was done using ClarityMax ECL (BUBR1 and CHK2) or Clarity ECL (all remaining proteins) reagents from BioRad and imaged on Azure600. All IP experiments were done in triplicate.

### Supplementary figures

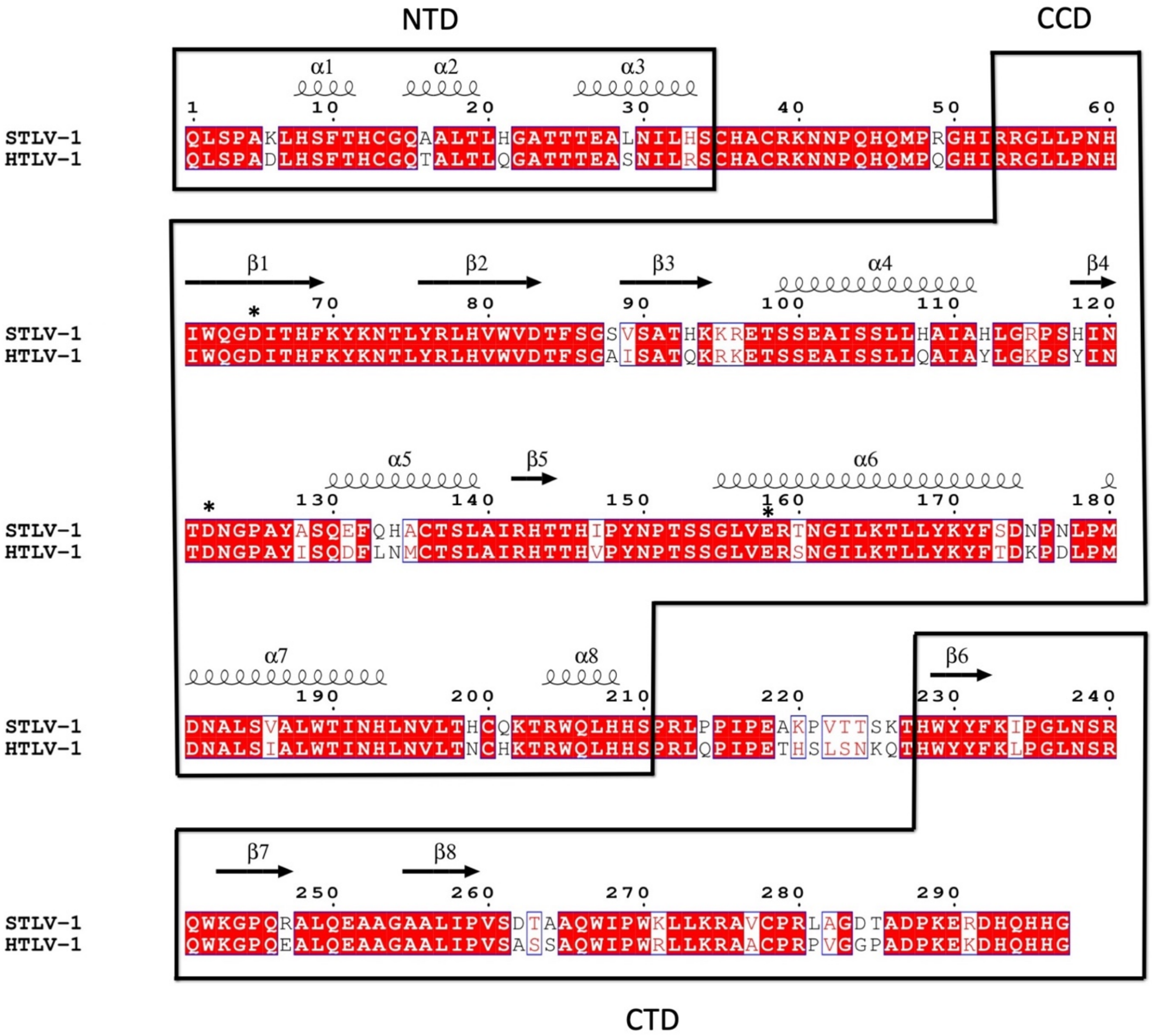

**Figure S1 | Sequence Alignment of STLV-1 MarB43 and HTLV-1 INs.** N-terminal (NTD), catalytic core (CCD) and C-terminal (CTD) domains are indicated with black boxes. Residue identity is indicated with red background, similarity with red lettering. Residues forming the catalytic triad DDE are indicated with asterisks. The STLV-1 MarB43 and HTLV-1 IN sequences exhibit 83% identity and 92% similarity. Alignment was conducted with CLUSTALW(30) and visualised in ESPRIPT (31).

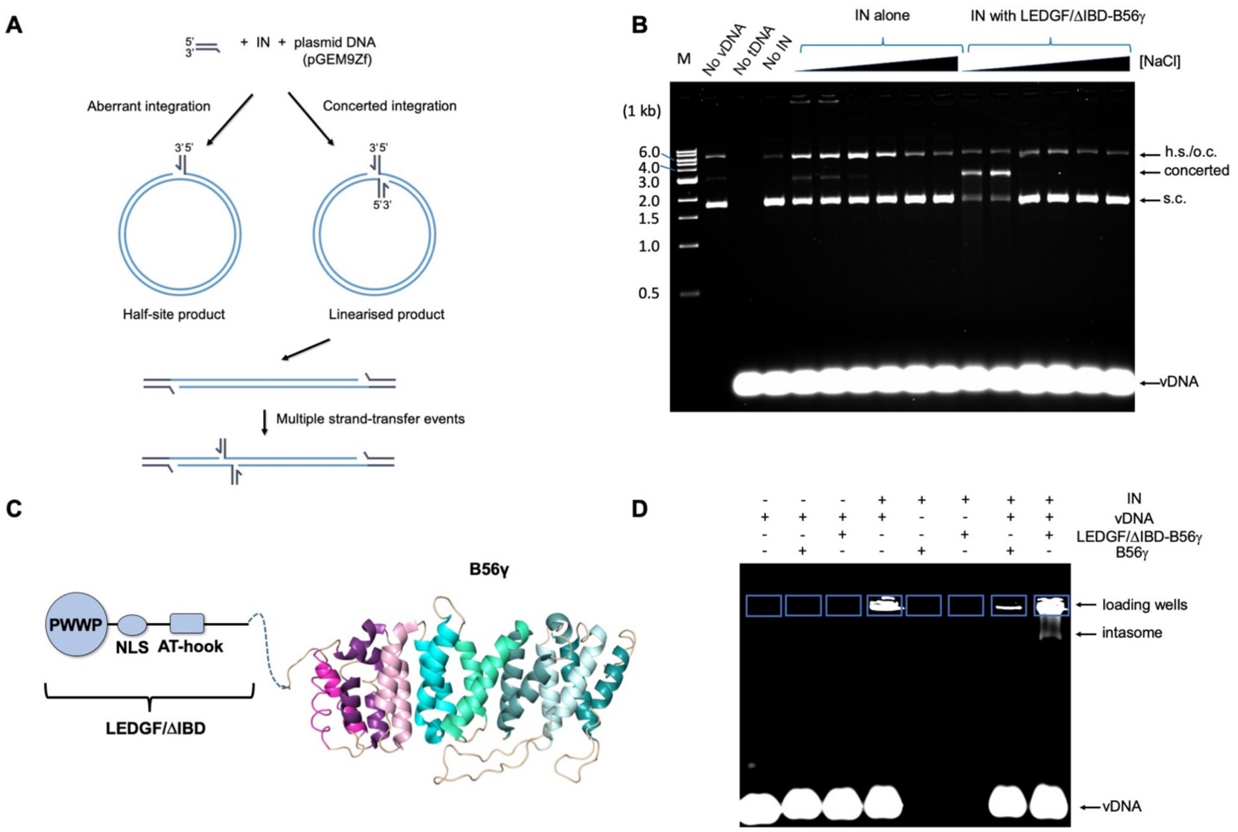

**Figure S2 | Optimisation of STLV-1 intasome activity and assembly. A**, IN strand-transfer activity can be measured by means of an assay where LTR mimics (vDNA) and target DNA (tDNA, a supercoiled plasmid) are mixed with recombinant STLV-1 IN. A linearised plasmid product corresponds to integration of both vDNA ends inserted. **B**, STLV-1 strand-transfer activity was measured at different NaCl concentrations during co-incubation with vDNA in presence or absence of LEDGF/ΔIBD-B56γ. The NaCl concentrations from left to right: 60 mM, 100 mM, 200 mM, 300 mM, 400 mM, 500 mM. M stands for the marker ladder (NEB, 1kb), h.s – half-site integration, o.c. – open circular, s.c. – supercoiled. Although residual activity of STLV-1 IN is visible without LEDGF/ΔIBD-B56γ, activity with the binding partner, as previously shown for B56γ on its own {Maertens, 2016 #46}, is considerably enhanced. NaCl concentrations above 100 mM have a negative impact on strand-transfer activity. **C**, Schematic of LEDGF/ΔIBD-B56γ structure. **D**, EMSA assay showing the dependence of successful STLV-1 intasome formation on the presence of LEDGF/ΔIBD-B56γ. B56γ alone is not sufficient to lead to stable intasome assembly. Atto680-labelled vDNA (30 bp) was used to visualise the DNA on a 3% low melting point agarose gel.

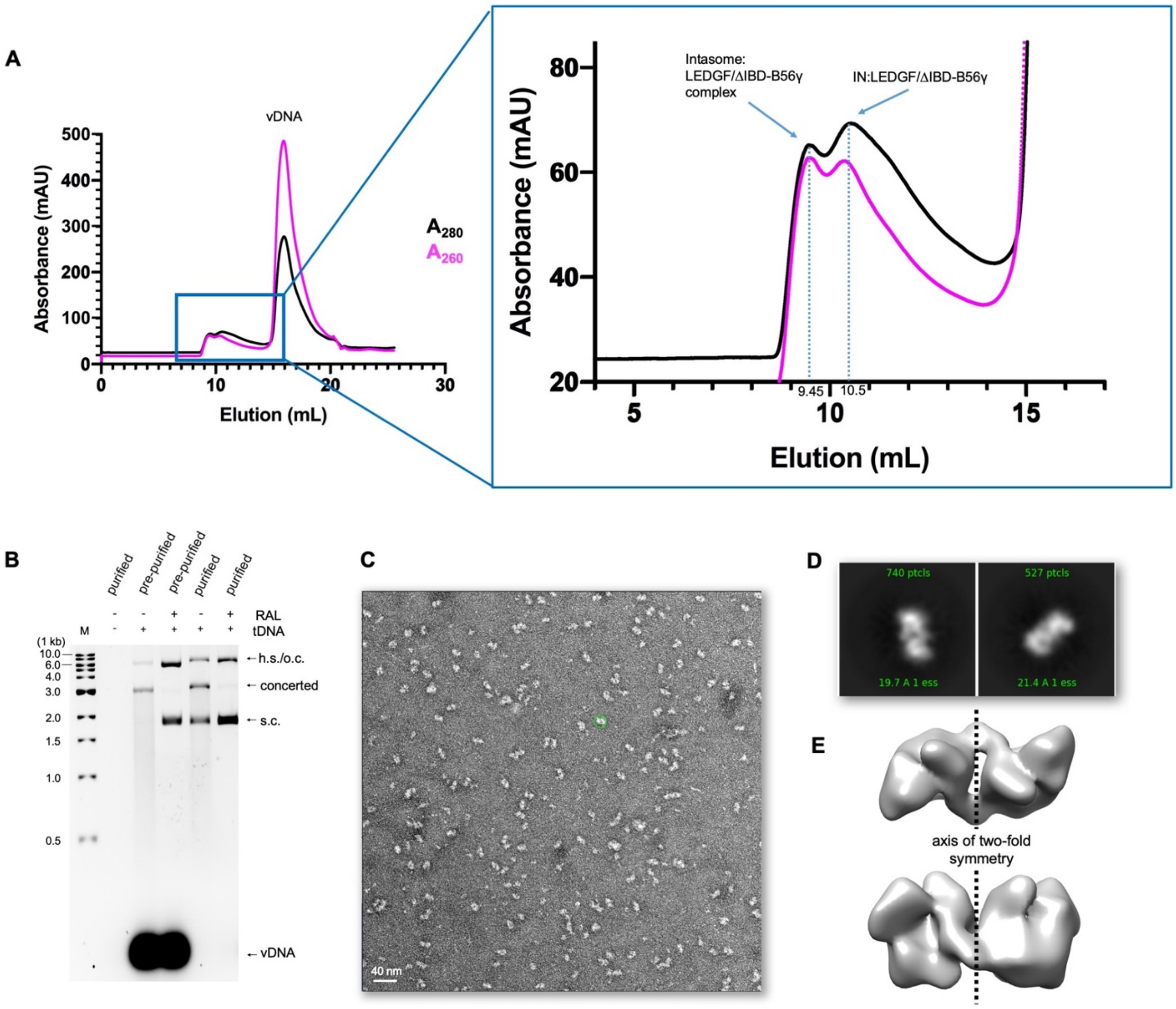

**Figure S3 | Assembly of STLV-1 intasomes for electron microscopy. A**, Purification of STLV-1 intasome by size exclusion chromatography (SEC). Two high-molecular weight peaks corresponding to the assembled STLV-1 intasome eluting at 9.45 mL and the smaller IN: LEDGF/ΔIBD-B56γ complex, devoid of vDNA, eluting at 10.5 mL. The black trace shows absorbance recorded at 280 nm, the purple trace absorbance at 260 nm. **B**, Analysis of strand-transfer activity associated with crude and SEC-purified intasome fractions in presence or absence of target DNA (tDNA) and the strand-transfer inhibitor raltegravir (RAL). Products were resolved on a 1.5% agarose gel, stained with ethidium bromide. M stands for the marker ladder (NEB, 1kb); h.s., half-site integration; o.c., open circular; s.c., supercoiled. **C**, Electron micrograph of the intasome SEC fraction negatively stained with uranyl acetate. An example of a particle is circled in green, measuring ~150 Å. **D**, Examples of 2D class averages of 22,000 negatively-stained intasome images. **E**, Orthogonal views of a 3D reconstruction obtained from 8,790 negatively-stained STLV-1 intasome images selected by reference-free 2D classification; two-fold symmetry axis is indicated.

A

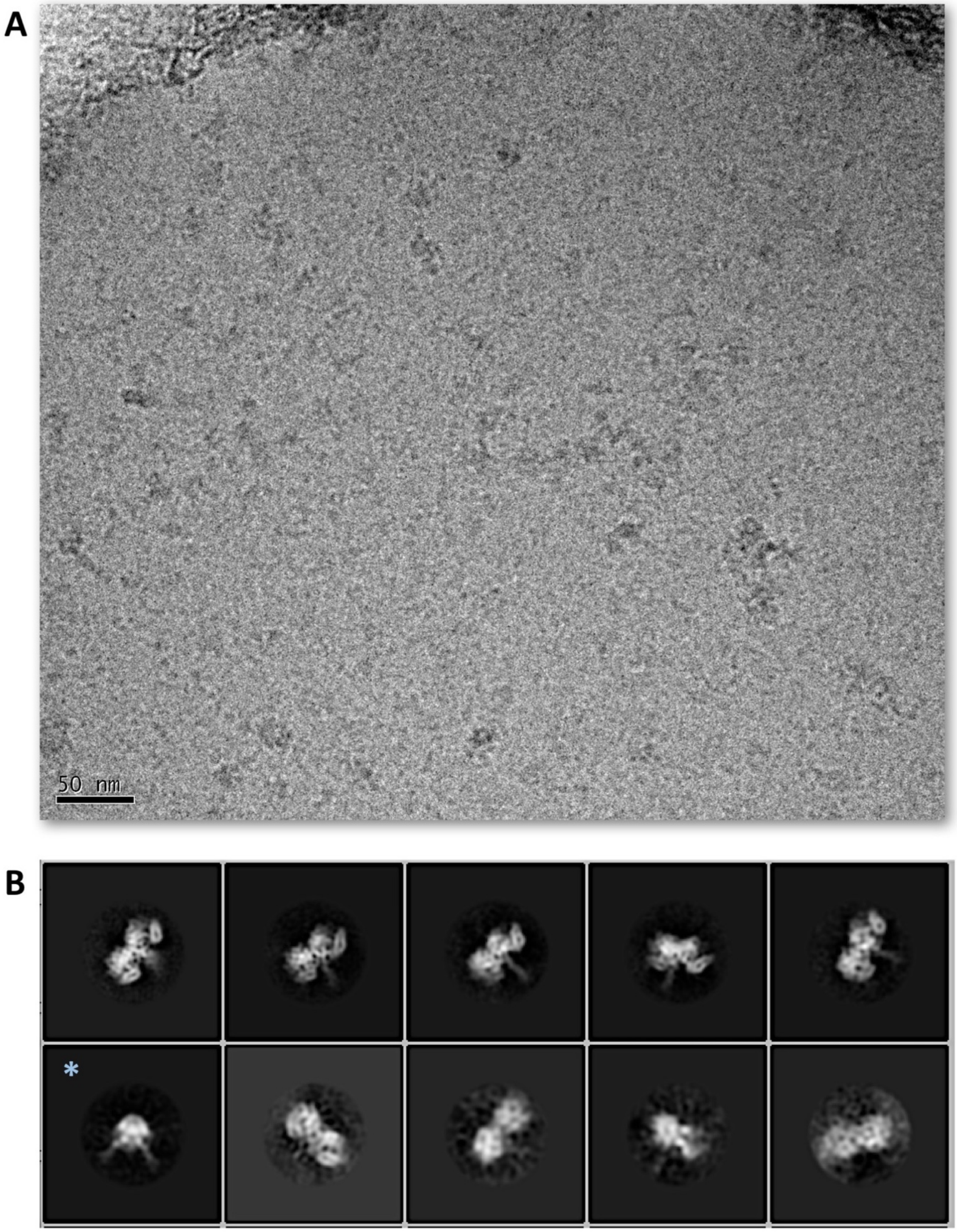

**Figure S4 | Initial CryoEM analysis of the STLV-1 intasome.** (A) An example of cryoEM micrograph showing STLV-1 intasome particles used for obtaining initial 2D classes (B). Most classes represent the "side" view of the intasome while the class marked with an asterisk (*) represent the "through" view.

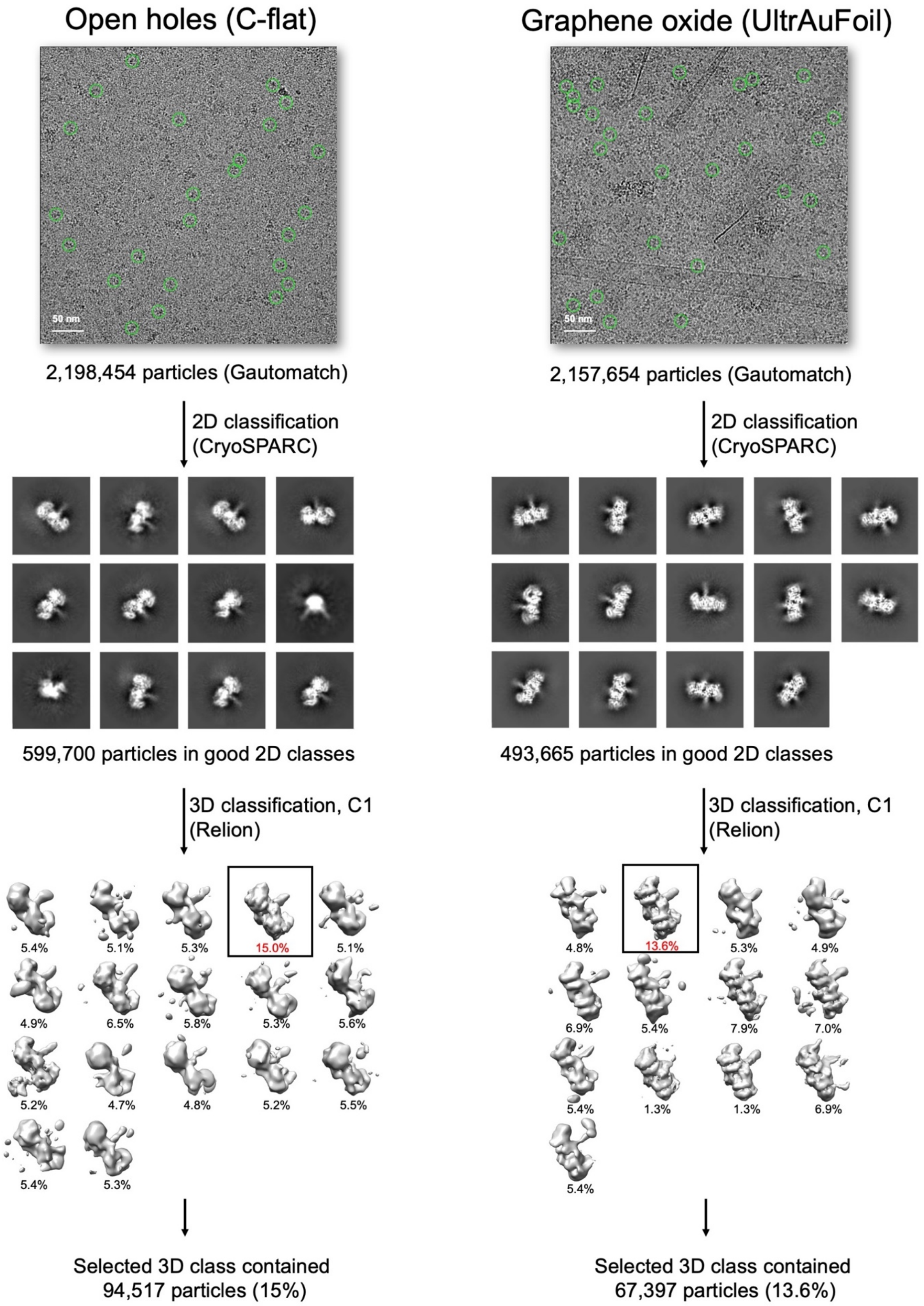
Open holes (C-flat)
Graphene oxide (UltrAuFoil)
50 nm
50 nm
2,198,454 particles (Gautomatch)
2,157,654 particles (Gautomatch)
2D classification (CryoSPARC)
2D classification (CryoSPARC)
599,700 particles in good 2D classes
493,665 particles in good 2D classes
3D classification, C1 (Relion)
3D classification, C1 (Relion)
5.4% 5.1% 5.3% 15.0% 5.1%
4.9% 6.5% 5.8% 5.3% 5.6%
5.2% 4.7% 4.8% 5.2% 5.5%
5.4% 5.3%
4.8% 13.6% 5.3% 4.9%
6.9% 5.4% 7.9% 7.0%
5.4% 1.3% 1.3% 6.9%
5.4%
Selected 3D class contained 94,517 particles (15%)
Selected 3D class contained 67,397 particles (13.6%)

**Figure S5 | Schematic of cryoEM image processing.** Open hole (OH) and graphene oxide (GO) datasets were processed separately prior to merging to alleviate severe anisotropy due to strong (but complementary) preferential particle orientations on OH and GO grids (Fig. S6, Table S5). Details are given in Materials and Methods.

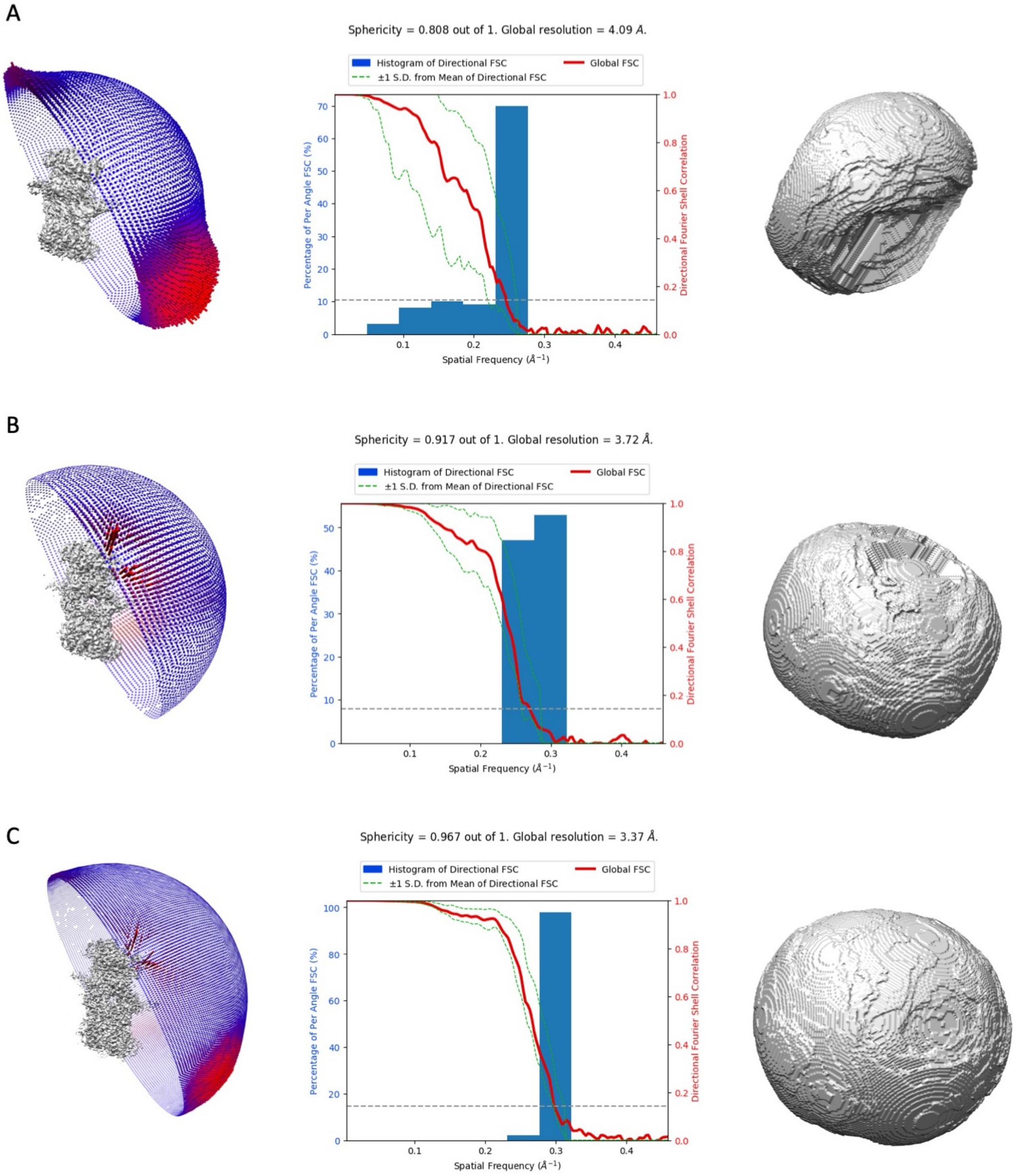

**Figure S6 | Orientational bias and anisotropy analysis.** Orientational bias was analysed for each of three datasets, corresponding to collections in open holes **(A)**, graphene oxide-coated grids **(B)**, and the merged final dataset **(C)**. The observed Euler angles (left), combined 2D FSC and 3D FSC histogram (middle) and binarized 3D FSC volumes (18) (right) are shown. Reconstructions from both OH and GO datasets suffer from considerable anisotropy, as indicated by presence of directions of low resolution (ranging between 12 and 4 Å for the map reconstructed from OH data) and non-spherical binarized 3D FSCs (A, B). Merging both datasets enriched the Euler angle distribution, significantly reducing the anisotropy and increasing the quantified sphericity score of the reconstruction (C).

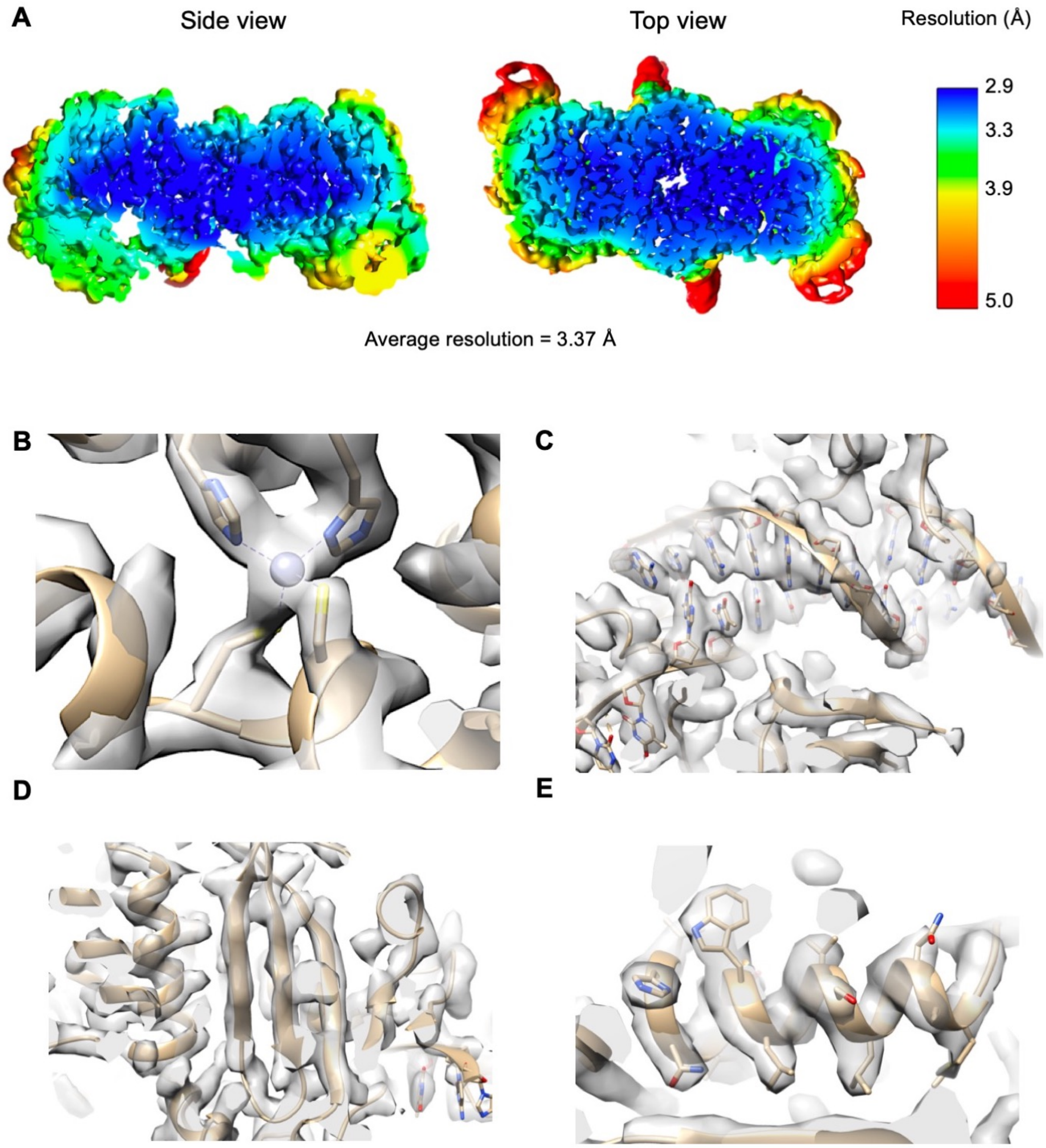

**Figure S7 | Analysis of the reconstructed STLV-1 map. A**, Local resolution of the reconstruction is mapped onto the density showing an average resolution of 3.37 Å, with the highest resolution (blue) of 2.9 Å around the core of the molecule (including the active site), and the lowest (red) in the outside helices of B56γ and terminal bases of vDNA. **B**, *Ab initio* building of the STLV-1 IN/NTD domain. Density for the $Zn^{2+}$ ion (blue sphere) is clearly resolved, together with the HHCC-binding motif. **C**, Building of the vDNA double helix. The double strand is then resolved in the intasome active site (left). **D**, Central β-sheet of the STLV-1 IN/CCD once fitted into the density. Clear separation of β-strands density is indicative of a sub-4 Å cryoEM reconstruction. **E**, High resolution of the intasome reconstruction allows for unambiguous placement of the majority of amino acid side chains; pictured is the IN/CCD helix $\alpha$7.

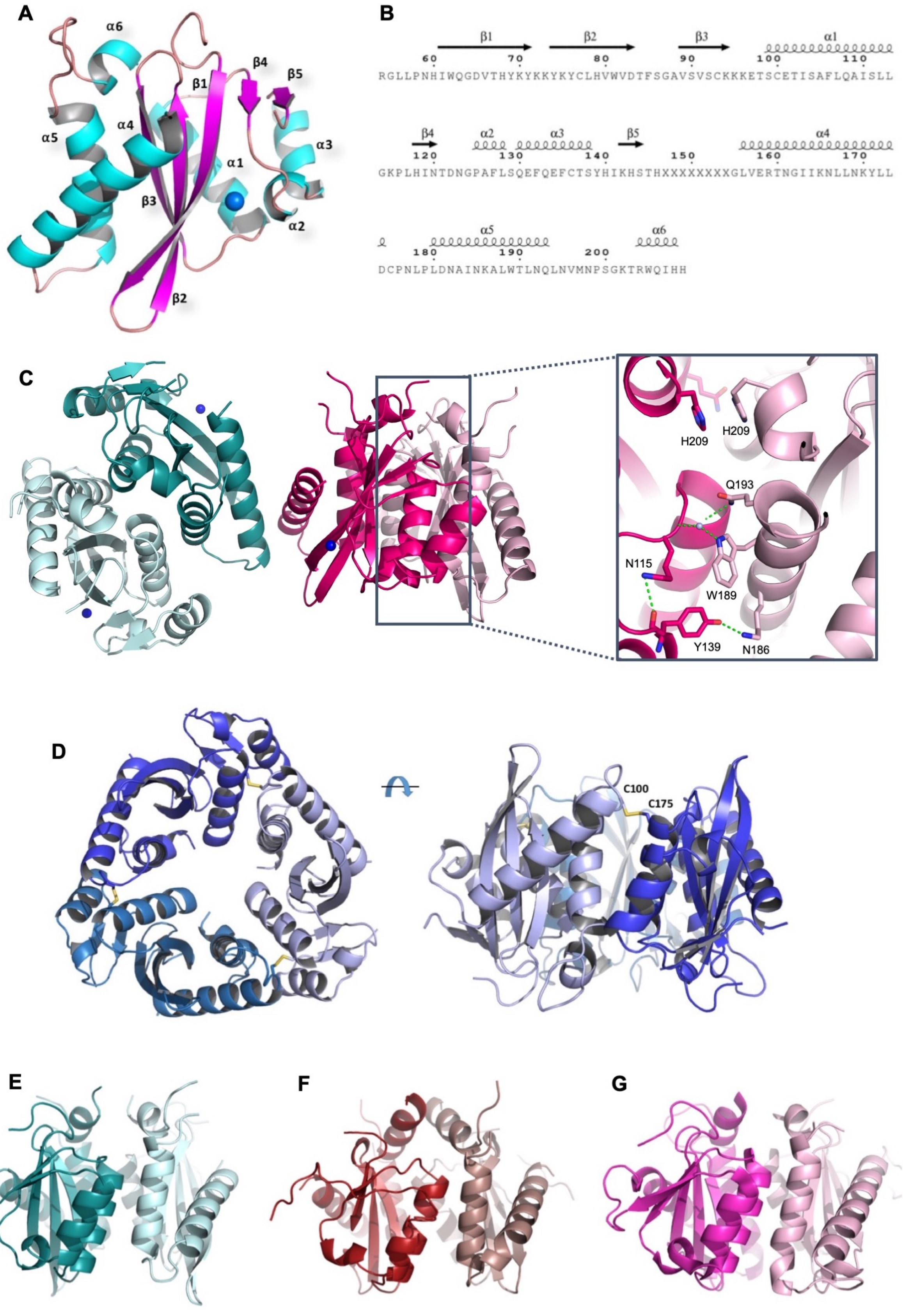
A
α6
α5
α4
β1
β4
β5
α3
α1
β3
α2
β2
B
β1
β2
β3
α1
60
70
80
90
100
110
RGLLPNHIWQGDVTHYKYKKYKYCLHVWVDTFSGAVSVSCKKKETSCETISAFLQAISLL
β4
α2
α3
β5
α4
120
130
140
150
160
170
GKPLHINTDNGPAFLSQEFQEFCTSYHIKHSTHXXXXXXXXGLVERTNGIIKNLLNKYLL
α5
α6
180
190
200
DCPNLPLDNAINKALWTLNQLNVMNPSGKTRWQIHH
C
H209
H209
Q193
N115
W189
Y139
N186
D
C100
C175
E
F
G

**Figure S8 | X-ray crystal structures of HTLV-2 IN/CCD. A**, Analysis of overall HTLV-2 IN/CCD structural features. Secondary structural features are shown as α-helices in cyan and β-strands in pink. The complexed magnesium atom is shown in blue. **B**, Amino acid sequence for HTLV-2 IN/CCD with the secondary structure topology overlaid. **C**, Crystal packing in the C121 space group reveals two distinct dimerisation interfaces, including the stable, canonical, form indicated with the grey rectangle. The interaction interface of the canonical dimer is shown in more detail in the inset (right). Residues important for the stabilisation of this interface are shown as sticks. Water molecules involved in these interactions are shown as light blue spheres and putative polar interactions as green dotted lines. **D,** Interestingly, we also crystallised an unusual trimeric assembly of IN/CCD formed between symmetry-related chains in the $P4_332$ space group. Although the interface buries 1,627 Å$^2$ of surface area (compared with 874 Å$^2$ buried by the canonical dimer), IN/CCD exists only as a dimer in solution (Figure S9). The disulphide bridges and the cysteines involved in trimerisation are indicated in yellow. **E-G**, Comparison of the highly structurally-conserved canonical dimer interfaces in IN/CCD structures of related retroviral genera. The panels represent CCD structures from (**E**) HTLV-2, (**F**) HIV-1 (PDB ID: 5KRT), and (**G**) RSV (PDB ID: 1C1A) IN.

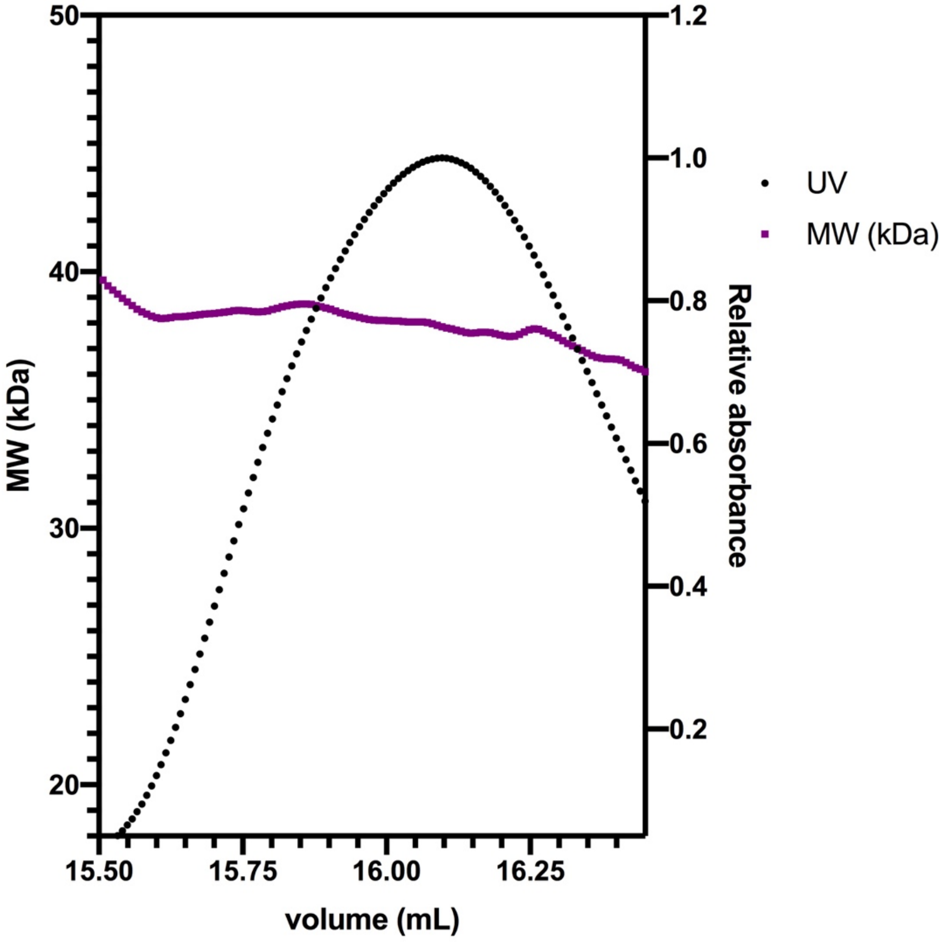

**Figure S9 | SEC-MALS analysis of HTLV-2 IN/CCD.** A purified sample of HTLV-2 IN/CCD (residues 53-221) was analysed in a SEC-MALS experiment to more precisely ascertain the molecular weight (MW) of its oligomeric state in solution. UV absorption was recorded and showed a single elution peak at 16.1 mL. The MW for the peak was calculated in the ASTRA software (Wyatt Technology, UK) from light scattering and differential refractive index. The calculated MW of 38.13 kDa would correspond to a dimer (MW of a monomer being 19.2 kDa) of IN/CCD.

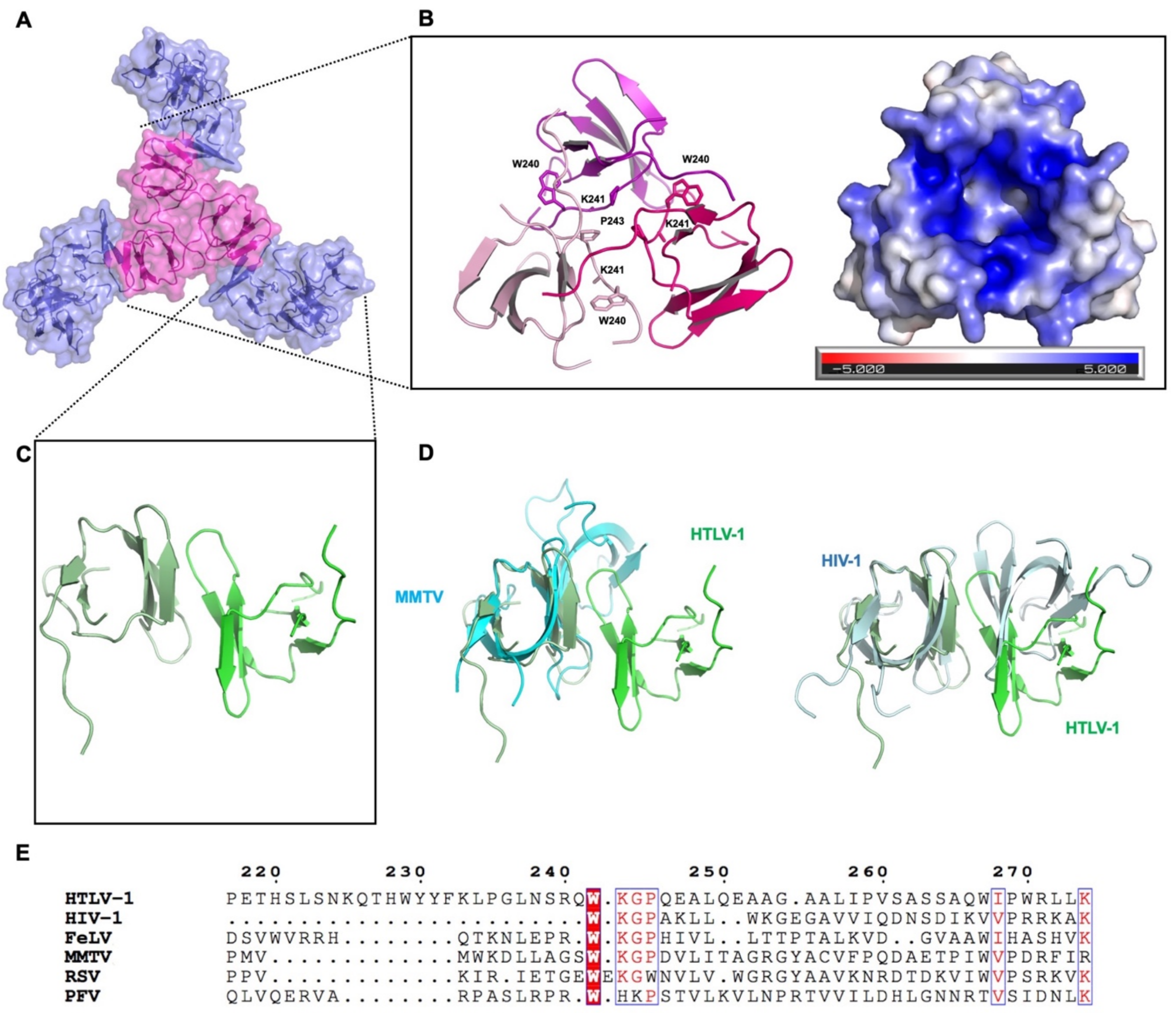

**Figure S10 | X-ray crystal structure of HTLV-1 IN/CTD. A**, A propeller-like trimer of IN/CTD dimers crystal packing of HTLV-1 IN/CTD leads to a canonical dimeric CTD structure (**C**) as well as a novel trimeric species (**B**). **B**, Curiously, the IN/CTD crystal packing creates a stable trimeric interface with a highly positively-charged core, involving residues Trp240-Pro244 – a motif known to be conserved across INs from all retroviral genera (see panel E) (32). Trimeric forms of IN have not been observed before and although likely not important for the catalytic function of IN, could be relevant to its other functions, such as vRNA binding (33). The stable trimeric interface harbours strong positive charge in its centre (right, as calculated with the APBS PyMol plugin), neutralised by water molecules. Red signifies negative charge, blue signifies positive charge. **C**, Dimeric packing of HTLV-1 IN/CTD. **D**, HTLV-1 IN/CTD dimers (green) compared to respective dimers observed in crystal and cryoEM structures of MMTV (cyan) and HIV-1 (light blue) INs. **E**, Alignment of IN/CTD sequences from different retroviral genera. The "WKGP motif" represents the most intra-genus and inter-genus conserved region of IN/CTD. Red background indicates absolute conservation, red font indicates incomplete conservation. Alignment was conducted with CLUSTALW (30) and visualised in ESPRIPT (31)

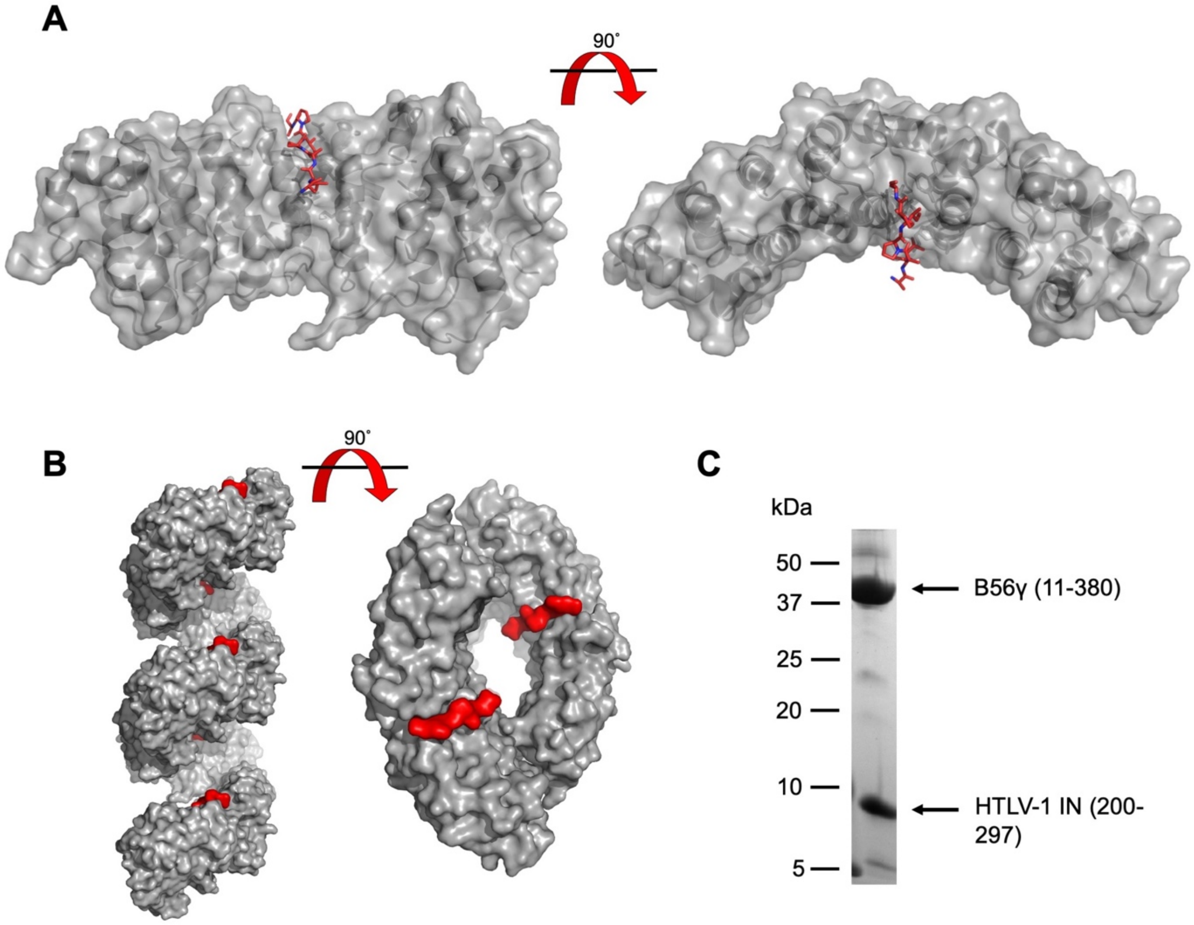

**Figure S11 | Structure of HTLV-1 IN (200-297) in complex with B56γ. A,** Overall crystal structure with B56γ (grey) seen in complex with the CCD-CTD linker region of HTLV-1 IN (red). **B**, Crystal packing creates helical arrangements of B56γ, creating large solvent channels. The resolved, SLiM-containing fragment of IN (red) is positioned facing the lumen of the helix. The remaining, unresolved part of HTLV-1 IN (200-297) is therefore likely to be disordered and contained in the solvent channels created by this crystal packing. **C,** The crystals used for diffraction were crushed, dissolved in running buffer and analysed on SDS-PAGE. This would indicate that despite the majority of HTLV-1 IN (200-297) density missing, the protein in its intact form is present in the crystal.

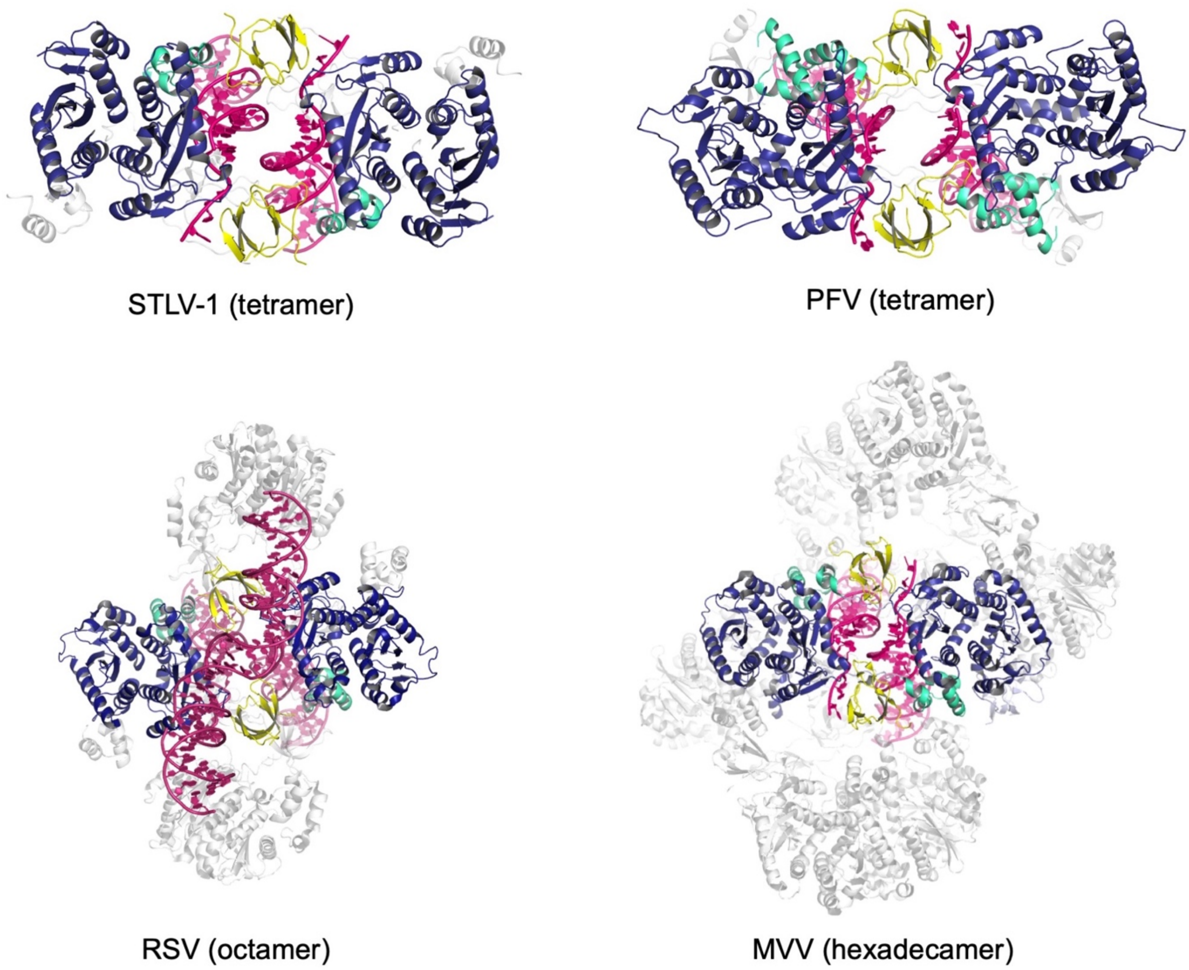

**Figure S12 | The STLV-1 intasome structure as compared to intasomes from other retroviral genera.** Structures of intasome complexes display surprising variability among the different retroviral genera. Although the oligomeric assemblies differ significantly, the conserved intasome core (CIC), coloured non-grey above, is largely unchanged. The structure of STLV-1 intasome is closest to that of PFV. B56γ was removed for clarity. IN/NTDs are shown in cyan, IN/CCDs in blue, IN/CTDs in yellow and vDNA in pink, and all residues outside of the CIC were coloured in semi-transparent grey. PDB accession codes are as follows: 6Z2Y (STLV-1), 3L2Q (PFV), 5EJK (RSV), 5M0Q (MVV).

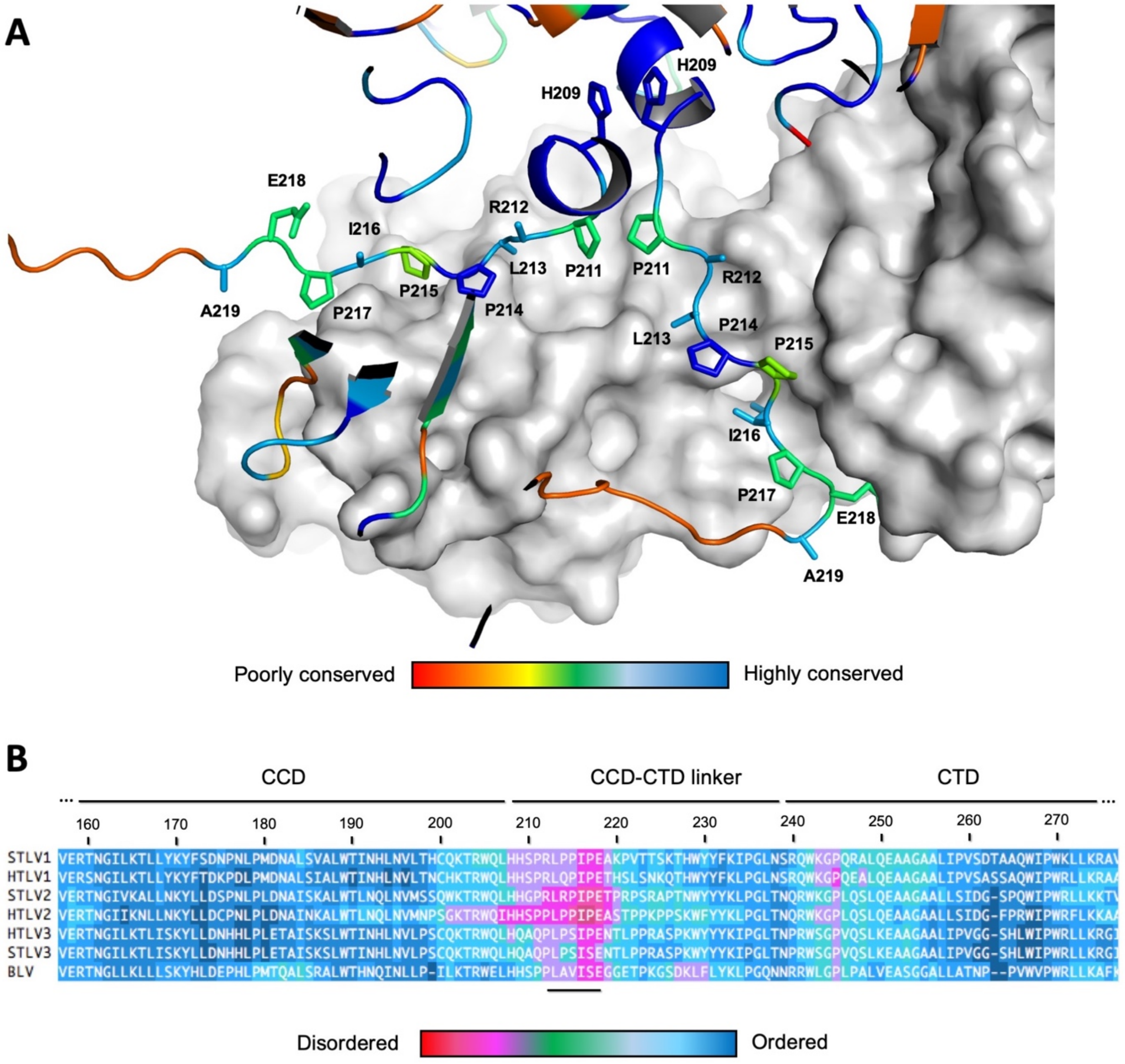

**Figure S13 | Sequence conservation and intrinsic disorder analysis of STLV-1 IN. A,** Sequence conservation between delta-retroviral IN linker regions was projected on the STLV-1 : B56γ intasome structure, showing high conservation in the SLiM region engaged in B56γ interactions in all resolved IN subunits. Residues $_{213}$LPPIPE$_{218}$ comprise the conserved LxxIxE interaction motif. Sequence conservation is coloured from blue (high conservation) to red (no conservation). Figure was prepared with Alebrijes 2.1 (https://github.com/mbarski/Alebrijes) **B**, Disorder prediction on aligned delta-retroviral INs shows that the SLiM-containing IN CCD-CTD linker is intrinsically disordered (LxxIxE motif, underlined). Conservation of disorder in this region among delta-retroviral INs is also clearly visible. Red indicates high disorder, dark blue - absence of disorder. Analysis was conducted with BASILIScan 1.3 (34).

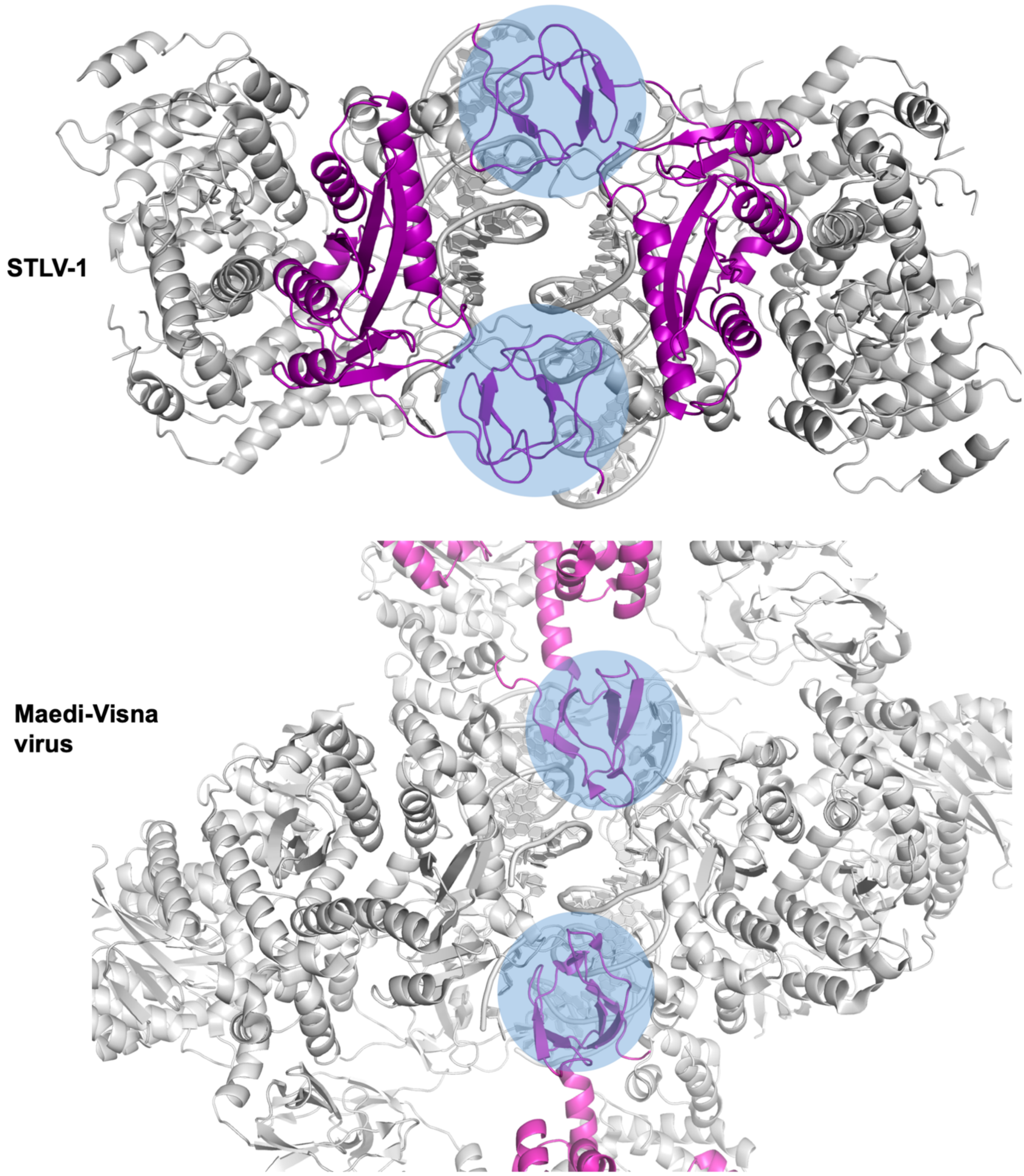

**Figure S14 | Positioning of the synaptic CTD domains in intasome structures.** The extended nature of the CCD-CTD linkers of STLV-1 IN allow for positioning of synaptic CTDs in *cis* (top), while INs with shorter, or coiled CCD-CTD linkers (for example MVV, bottom) require positioning of the synaptic CTD in *trans*, engaging flanking IN subunits and leading to a larger oligomeric state of the intasome. For clarity the B56γ host factor was removed from the STLV-1 intasome structure (top). PDB accession codes: 6Z2Y (STLV-1), and 5M0Q (MVV).

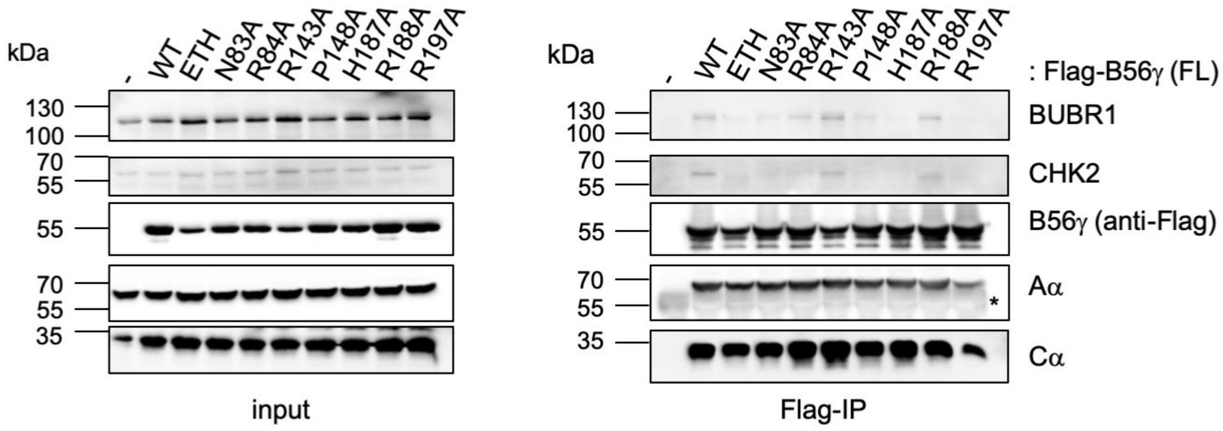

**Figure S15 | Co-immunoprecipitation (IP) of endogenous PP2A-B56γ substrates.** Extracts from 293T cells transiently transfected with Flag-tagged full-length B56γ WT or mutant versions were immunoprecipitated with anti-Flag antibodies. Binding to endogenous substrates (BUBR1 and CHK2) as well as formation of the holo-enzyme was investigated by western blot. Antibodies used are indicated to the right of the blots, Mw markers are indicated to the left. Mutants are shown on top of the blots. As a negative control empty vector was transfected into 293T cells (-). Input and Flag-IP samples are shown in the left and the right panel, respectively. *indicates cross-reaction of the secondary anti-rat antibody with the mouse anti-Flag antibody used during IP. This is a representative of three biological replicates.

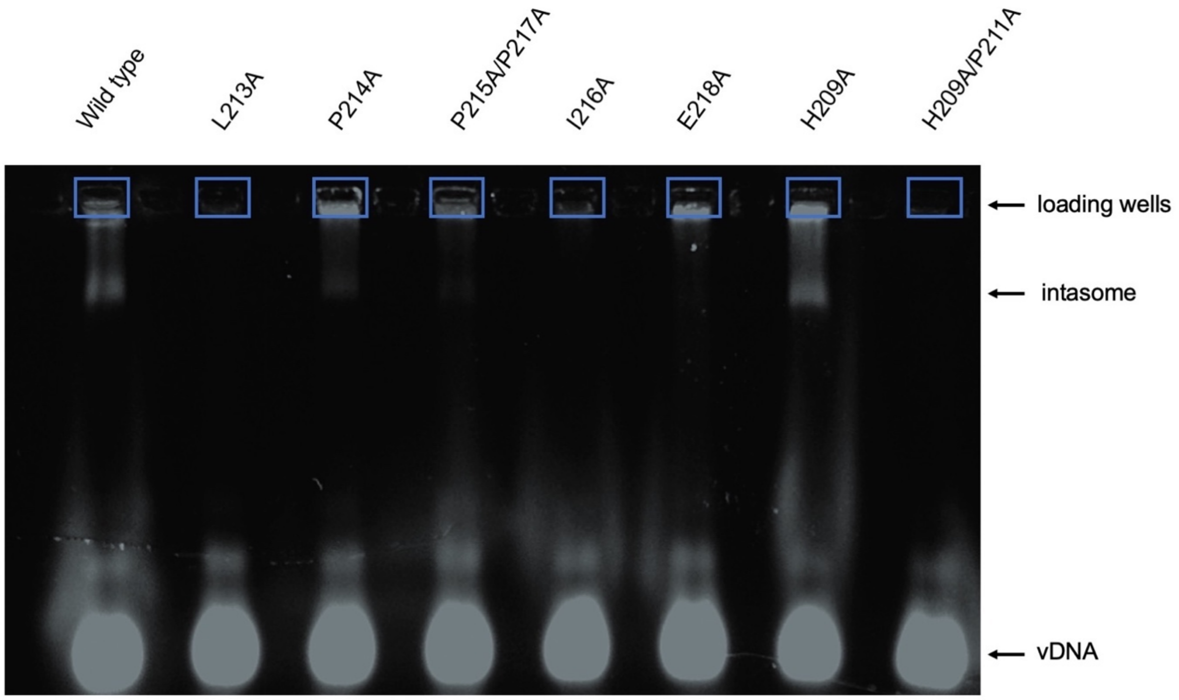

**Figure S16 | Electrophoretic mobility shift assays analysis of STLV-1 IN mutants.** Representative image of EMSA used for quantification shown in Figure 2. As seen in Figure S2, LEDGF/ΔIBD-B56γ is crucial for intasome assembly *in vitro*. Mutations of IN residues seen involved in binding to B56γ in the cryoEM structure affect assembly of the intasome. Atto680-labelled vDNA (30 bp) was used to visualise the DNA on a 3% low melting point agarose gel. Experiment was conducted in triplicate for quantification shown in Figure 2.

### Supplementary tables

**Table S1**. Primers used to clone constructs for recombinant protein expression

| Primer name | Primer sequence |
|---|---|
| GM110 | CCGGGTCGACTCAGCCGTGGTGCTGGTGGTC |
| GM130 | GGCCCAATTGCGCCGGGGCCTCTTGCC |
| GM142 | GCTTGAATTCATGGTGGTGGATGCGGC |
| GNM283 | GGCCGTCGACTCAGGTCTTTGAGTTGCGG |
| GNM367 | GGCCGTCGACCTAGCGGCCGTCCTGGG |
| GNM378 | GGCCGAATTCTGCCACAAGACCCGGTGGC |
| GNM380 | GGCCACCGGTATGTTGACATGTAATAAAGCGGGC |
| GNM641 | CCGGGGATCCATGACTCGCGATTTCAAACC |
| GNM747 | GGCCGAATTCCAACTTTCTCCTGCCAAACTACATAGTTTTACTC |
| GNM748 | GGCCCTCGAGTTACCCATGGTGTTGGTGGTCTCTTTC |
| GNM750 | CTGGGTAAATAGGCTCTGTGATCACATTCCGATTAGCGGCGATATATGCTACCATTTCAC |
| GNM751 | CAGCTTGTTTATGAATTTTTCTTAGCATTTTTAGAGTCTCCAGATTTCCAACCTAATATAGCG |
| GNM752 | CGCTATATTAGGTTGGAAATCTGGAGACTCTAAAAATGCTAAGAAAAATTCATAAACAAGCTG |
| GNM769 | GGCCGAATTCCAGCTGAGTCCGGCAAAACTGCATAG |
| GNM769 | GGCCGAATTCCAGCTGAGTCCGGCAAAACTGCATAG |
| GNM770 | GGCCCTCGAGTTAACCATGATGCTGATGATCACGTTC |
| GNM771 | GCTGCATCATAGTCCGCGTGCGCCTCCGATTCCGGAAGCAAAACC |
| GNM772 | GGTTTTGCTTCCGGAATCGGAGGCGCACGCGGACTATGATGCAGC |
| GNM773 | GCATCATAGTCCGCGTCTGGCTCCGATTCCGGAAGCAAAACC |
| GNM774 | GGTTTTGCTTCCGGAATCGGAGCCAGACGCGGACTATGATGC |
| GNM775 | CATCATAGTCCGCGTCTGCCTGCGATTGCGGAAGCAAAACCGGTTACCACC |
| GNM776 | GGTGGTAACCGGTTTTGCTTCCGCAATCGCAGGCAGACGCGGACTATGATG |
| GNM777 | CATAGTCCGCGTCTGCCTCCGGCTCCGGAAGCAAAACCGGTTACC |
| GNM778 | GGTAACCGGTTTTGCTTCCGGAGCCGGAGGCAGACGCGGACTATG |
| GNM779 | GTCCGCGTCTGCCTCCGATTCCGGCAGCAAAACCGGTTACCACCAG |
| GNM780 | CTGGTGGTAACCGGTTTTGCTGCCGGAATCGGAGGCAGACGCGGAC |
| GNM824 | GAATCGGAGGCAGACGCGGACTAGCATGCAGCTGCCAACGGGTTTTC |
| GNM826 | CCGGAATCGGAGGCAGACGCGCACTAGCATGCAGCTGCCAACGGGTTTTC |
| GNM829 | GCTGCATGCTAGTCCGCGTCTGCCTCCGATTC |

| | |
|---|---|
| GNM830 | GCTGCATGCTAGTGCGCGTCTGCCTCCGATTCCGG |
| GNM833 | ATGGTAGAATATATCACCCATAATGCGAATGTGATCACAGAGCCTATTTACCCAGAAG |
| GNM834 | CTTCTGGGTAAATAGGCTCTGTGATCACATTCGCATTATGGGTGATATATTCTACCAT |
| GNM837 | GTTTATGAATTTTTCTTAAGATTTTTAGAGTCTGCAGATTTCCAACCTAATATAGCG |
| GNM838 | CGCTATATTAGGTTGGAAATCTGCAGACTCTAAAAATCTTAAGAAAAATTCATAAAC |
| GNM844 | GGTAGAATATATCACCCATGCTCGGAATGTGATCACAGAGC |
| GNM845 | GCTCTGTGATCACATTCCGAGCATGGGTGATATATTCTACC |
| JM1 | GATCCCATCACCATCACCACCATGGCAGCGGCCTGGAAGTGCTGTTTCAAGGCCCGG |
| JM2 | AATTCCGGGCCTTGAAACAGCACTTCCAGGCCGCTGCCATGGTGGTGATGGTGATGG |
| JM3 | GGCCCTCGAGCTAGCGGCCGTCCTGGG |
| MB054 | TATGTCGACTCAAGCAGGTCCGCCC |
| MB094 | TATGTCGACTCAGGTAGAGGCTTCAG |
| MB114 | ATAGAATTCCACTGGTACTACTTCAAG |

**Table S2**. Primers used for STLV-1 IN strand-transfer assays

| **Name** | **Sequence** |
|---|---|
| Mar_U5_S30UP | TCTCTCCGGGAGAGAAGCGCCAAACACA |
| Mar_U5_S30B | ACTGTGTTTGGCGCTTCTCTCCCGGAGAGA |

**Table S3.** Data collection and refinement statistics of HTLV-2 IN CCD crystal structures.

| | IN CCD-$Mg^{2+}$ | IN CCD-$Mg^{2}$ | IN CCD-$Ca^{2+}$ |
|---|---|---|---|
| **Data collection** | | | |
| Space group | $P4_332$ | C121 | $P4_332$ |
| Cell dimensions | | | |
| *a*, *b*, *c* (Å) | 115.9, 115.9, 115.9 | 185.3, 89.2, 65.5 | 112.1, 112.1, 112.1 |
| α, β, γ (°) | 90, 90, 90 | 90, 103, 90 | 90, 90, 90 |
| Resolution (Å) | 40.98 - 2.29 (2.37 - 2.29) | 58.54 – 2.45 (2.55 – 2.45) | 50.12 – 2.40 (2.49 – 2.40) |
| $R_{merge}$ (%) | 2.3 (29) | 24.2 (38.3) | 2.8 (31) |
| $<I / \sigma(I)>$ | 24.2 (2.5) | 3.0 (1.2) | 13.8 (1.8) |
| Completeness (%) | 99.9 (99.9) | 100 (100) | 100 (100) |
| Redundancy | 1.8 (1.9) | 3.9 (3.9) | 1.8 (1.9) |
| **Refinement** | | | |
| Resolution (Å) | 2.29 | 2.45 | 2.40 |
| No. reflections | 22,653 (2,228) | 149,376 (16,967) | 17,832 (1,872) |
| $R_{work}$ / $R_{free}$ | 19.66/23.91 | 24.99/27.64 | 24.71/28.19 |
| No. atoms | | | |
| Protein | 1,195 | 4,640 | 1,190 |
| Ligand/ion | 1 | 4 | 1 |
| Water | 59 | 100 | 28 |
| *B*-factors (Å$^2$) | | | |
| Protein | 38.95 | 70.57 | 55.01 |
| Ligand/ion | 53.35 | 72.75 | 69.99 |
| Water | 64.19 | 81.82 | 84.59 |
| R.m.s. deviations | | | |
| Bond lengths (Å) | 0.017 | 0.011 | 0.005 |
| Bond angles (°) | 2.047 | 1.42 | 0.860 |

The values in parenthesis refer to the highest resolution shell.

[a] $R_{merge} = \frac{\sum_{hkl}\sum_i |I(hkl;i) - <I(hkl)>|}{\sum_{hkl}\sum_i I(hkl;i)}$, where I(hkl; i) is the intensity of an individual measurement of a reflection and <I(hkl)> is the average intensity of that reflection.

[b] $R_{work} = \frac{\sum_{hkl} F_0 - F_c}{\sum_{hkl} F_0}$, where $F_o$ and $F_c$ are the observed and calculated structure factors, respectively.

[c]$R_{free}$ is $R_{work}$ with 5% of the observed reflections removed before refinement.

**Table S4.** Data collection and refinement statistics of HTLV-1 IN CTD and HTLV-1 IN(200-297) : B56γ crystal structures.

| | **HTLV-1 IN CTD** | **HTLV-1 IN(200-297)-B56γ** |
|---|---|---|
| **Data collection** | | |
| Space group | $I4_132$ | $P4_32_12$ |
| Cell dimensions | | |
| *a*, *b*, *c* (Å) | 97.41, 97.41, 97.41 | 58.77, 58.77, 321.15 |
| α, β, γ (°) | 90, 90, 90 | 90, 90, 90 |
| Resolution (Å) | 39.80 – 1.80 (1.85 – 1.80) | 80.29 – 3.16 (3.38 – 3.16) |
| $R_{merge}$ (%) | 7.7 (71) | 16.7 (48.9) |
| $<I / (\sigma I)>$ | 7.4 (1.8) | 3.3 (1.5) |
| Completeness (%) | 100 (100) | 96.7 (94.4) |
| Redundancy | 5.4 (5.6) | 1.4 (1.2) |
| **Refinement** | | |
| Resolution (Å) | 1.8 | 3.16 |
| No. reflections | 41,483 (2,513) | 14,094 (2,025) |
| $R_{work}$ / $R_{free}$ | 17.71/24.64 | 22.7/29.4 |
| No. atoms | | |
| Protein | 840 | 2,674 |
| Ligand/ion | 0 | 0 |
| Water | 57 | 46 |
| *B*-factors | | |
| Protein | 55.00 | 53.90/79.71 |
| Ligand/ion | N/A | N/A |
| Water | 71.53 | 40.47 |
| R.m.s. deviations | | |
| Bond lengths (Å) | 0.012 | 0.0216 |
| Bond angles (°) | 1.861 | 1.98 |

The values in parenthesis refer to the highest resolution shell.

[a] $R_{merge} = \frac{\sum_{hkl}\sum_i |I(hkl;i) - <I(hkl)>|}{\sum_{hkl}\sum_i I(hkl;i)}$, where I(hkl; i) is the intensity of an individual measurement of a reflection and <I(hkl)> is the average intensity of that reflection.

[b] $R_{work} = \frac{\sum_{hkl} F_0 - F_c}{\sum_{hkl} F_0}$, where $F_o$ and $F_c$ are the observed and calculated structure factors, respectively.

[c]$R_{free}$ is $R_{work}$ with 5% of the observed reflections removed before refinement.

**Table S5.** CryoEM reconstruction and STLV-1 intasome refinement statistics.

| | STLV-1 intasome | |
|---|---|---|
| **Data collection** | OH subset | GO subset |
| Voltage (keV) | 300 | 300 |
| Cumulative exposure ($e^{-}/Å^{2}$) | 34 | 34 |
| Number of frames per movie | 30 | 30 |
| Defocus range (µm) | -1.6 to -3.6 | -1.6 to -3.6 |
| Pixel size (Å) | 1.09 | 1.09 |
| **Particle classification** | | |
| Initial particle images (no.) | 2,198,454 | 2,157,654 |
| Particles after 2D classification | 599,700 | 493,665 |
| Particles after 3D classification | 94,517 | 67,404 |
| **Merged 3D reconstruction** | | |
| Particles used | 161,914 | |
| Symmetry imposed | C2 | |
| Software for reconstruction | Relion-3.1 | |
| Overall resolution (Å)[a] | 3.37 | |
| Software for density modification and map sharpening | Resolve(Phenix-1.18-3845) and Relion-3.1 | |
| EMDB accession code | EMD-11052 | |
| **Model Refinement** | | |
| Final model (PDB accession code) | 6Z2Y | |
| Model composition | | |
| Non-hydrogen atoms | 14,998 | |
| Protein residues | 1,664 | |
| DNA | 80 | |
| $Zn^{2+}$ | 4 | |
| *B* factors ($Å^{2}$) | | |
| Protein | 31.31 | |
| DNA | 49.12 | |
| $Zn^{2+}$ | 38.96 | |
| **Model validation** | | |
| MolProbity score | 1.42 | |
| Clashscore | 3.63 | |
| EMringer score | 2.88 | |
| Poor rotamers (%) | 0.69 | |
| R.m.s. deviations | | |
| Bond lengths (Å) | 0.004 | |
| Bond angles (°) | 0.571 | |
| Ramachandran plot: | | |
| Favored (%) | 96.10 | |
| Allowed (%) | 100 | |
| Disallowed (%) | 0 | |

[a]Based on the FSC of 0.143 between half-sets

**Table S6.** Comparison of CCD-CTD linker lengths and respective oligomeric state of vDNA-complexed and free IN of retroviral genera for which intasome structures are available ordered according to linker length.

| Genus | Virus | CCD-CTD linker length | Intasome oligomer | Uncomplexed IN | Reference |
|---|---|---|---|---|---|
| Spumavirus | PFV | 50 | Tetramer | Monomer | (35) |
| Deltaretrovirus | STLV-1 | 19 | Tetramer | Dimer | This study |
| Lentivirus | MVV | 20 | Hexadecamer | Octamer | (36) |
| Betaretrovirus | MMTV | 8 | Octamer | Dimer | (37) |
| Alpharetrovirus | RSV | 8 | Octamer | Dimer | (38) |
